# Stable mammalian serum albumins designed for bacterial expression

**DOI:** 10.1101/2023.03.28.534334

**Authors:** Olga Khersonsky, Moshe Goldsmith, Irina Zaretsky, Shelly Hamer-Rogotner, Orly Dym, Tamar Unger, Meital Yona, Yael Fridmann-Sirkis, Sarel J. Fleishman

## Abstract

Albumin is the most abundant protein in the blood serum of mammals and has essential carrier and physiological roles. Albumins are also used in a wide variety of molecular and cellular experiments and in the cultivated meat industry. Despite their importance, however, albumins are challenging for heterologous expression in microbial hosts, likely due to 17 conserved intramolecular disulfide bonds. Therefore, albumins used in research and biotechnological applications either derive from animal serum, despite severe ethical and reproducibility concerns, or from recombinant expression in yeast or rice. We use the PROSS algorithm to stabilize human and bovine serum albumins, finding that all are highly expressed in *E. coli*. Design accuracy is verified by crystallographic analysis of a human albumin variant with 16 mutations. This albumin variant exhibits ligand binding properties similar to those of the wild type. Remarkably, a design with 73 mutations relative to human albumin exhibits over 40°C improved stability and is stable beyond the boiling point of water. Our results suggest that proteins with many disulfide bridges have the potential to exhibit extreme stability when subjected to design. The designed albumins may be used to make economical, reproducible, and animal-free reagents for molecular and cell biology. They also open the way to high-throughput screening to study and enhance albumin carrier properties.

**Graphical abstract:** 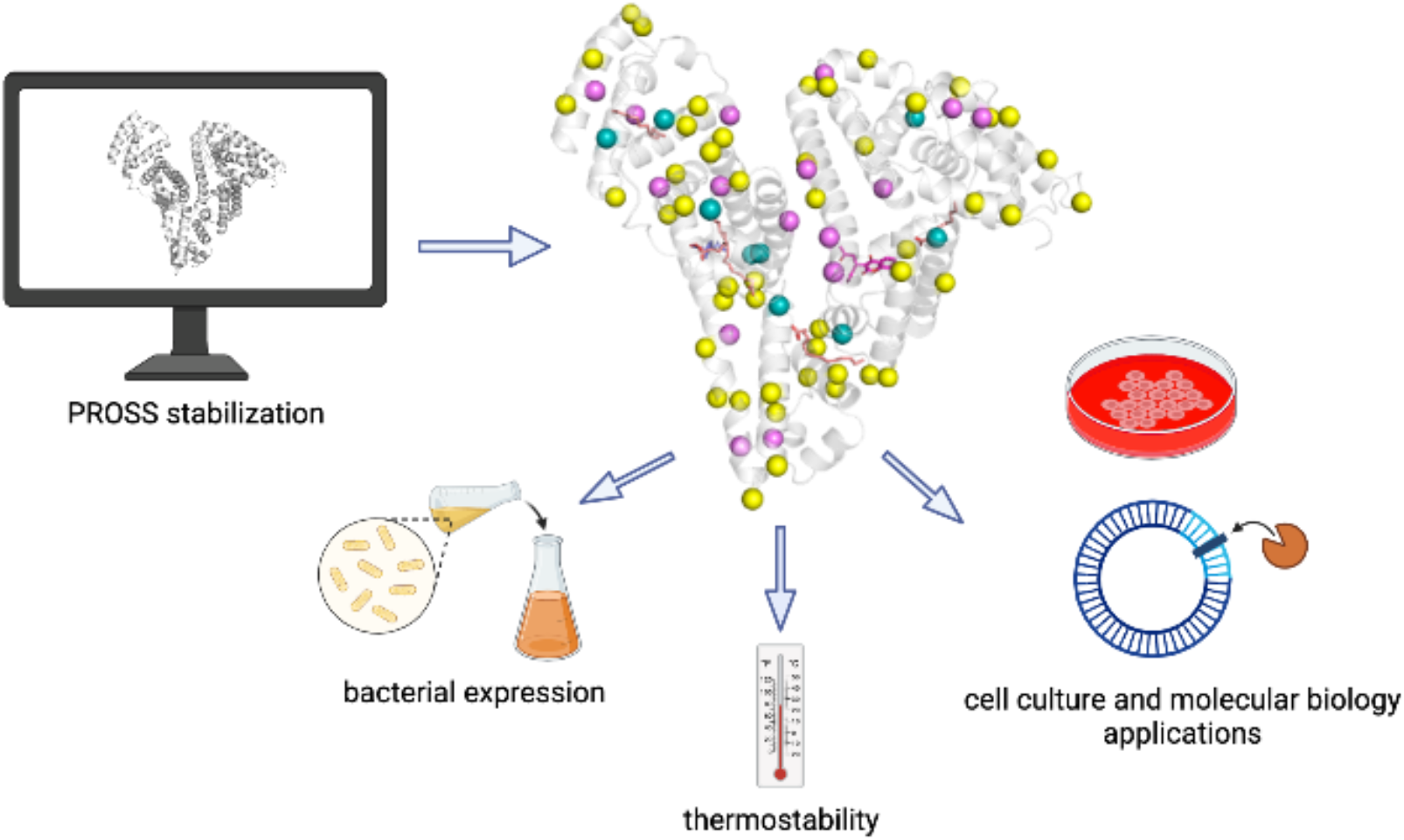

**Highlights:**

- Computational design stabilized human and bovine serum albumins
- Designs express solubly in *E. coli* and exhibit up to 40 °C increased thermostability
- Some designs exhibit identical ligand binding properties
- Crystal structure confirms design accuracy
- Designs can be used in cell culture and *in vitro* applications

## Introduction

Serum albumin is a monomeric, non-glycosylated, 67 kDa transport protein that is highly abundant in mammalian plasma (35-55 g/L) and has diverse physiological roles. It folds into a canonical heart-shaped structure comprising three helical domains with 17 conserved disulfide bridges (1)(**Fig. 1A**). Serum albumin is the main blood carrier for metabolites, hormones, drugs, and cations. It has seven binding sites for fatty acids and at least three binding sites for small molecules. Crystallographic analyses reveal two major conformations: compact and myristate-bound (**Fig. 1B**). Besides ligand binding, serum albumin has some catalytic properties and has a role in regulation of plasma colloid oncotic pressure(2–5).

**Fig. 1.**
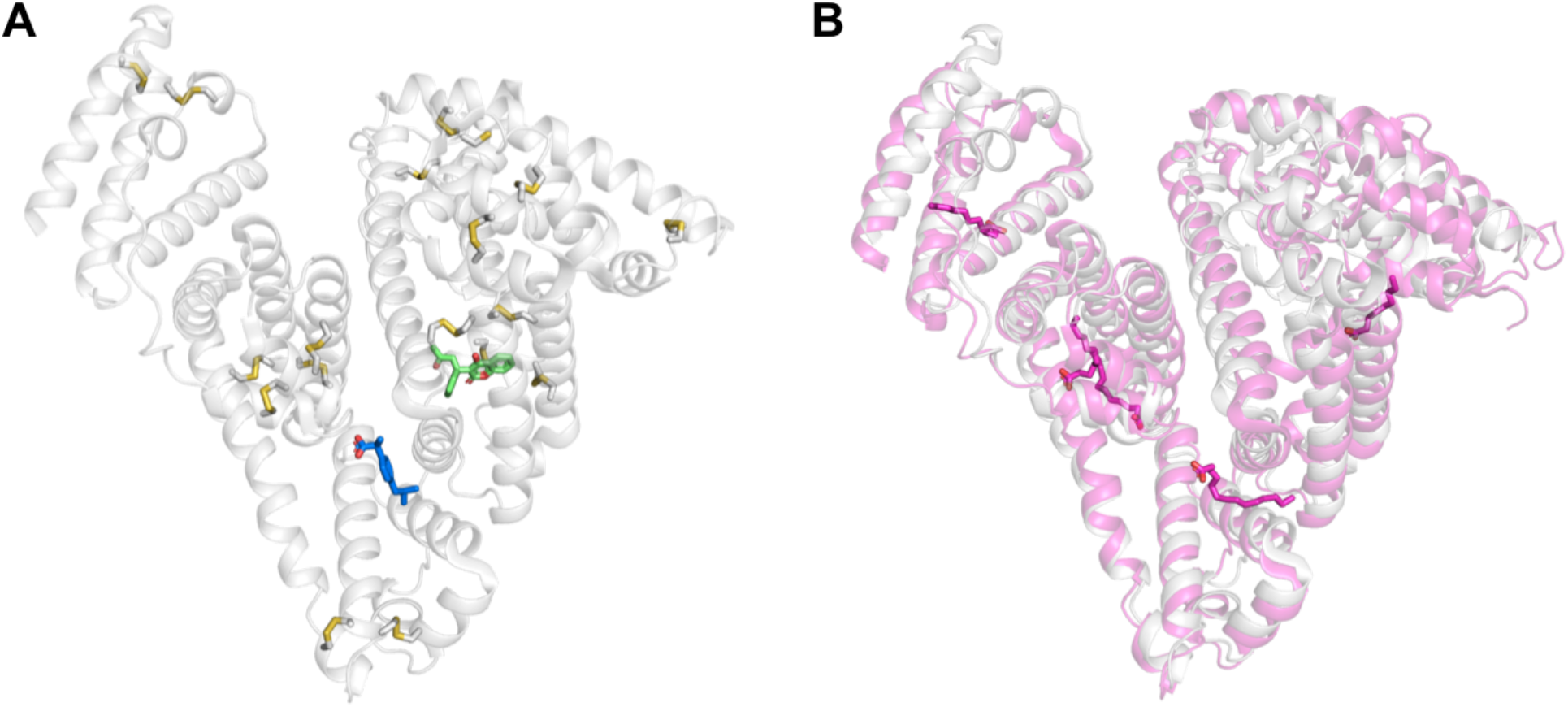
**(A)** The compact conformation of human serum albumin without myristate (PDB entry 2bxg). Disulfide bonds shown in gray sticks, ibuprofen in blue sticks, and warfarin in green sticks (overlaid from PDB entry 2bxd). **(B)** Human serum albumin structure, overlay of compact conformation (gray) with the open conformation (magenta, PDB entry 2bxi), with myristate molecules shown as pink sticks.

In addition to their physiological importance, human and bovine serum albumins (HSA and BSA, respectively) have many biochemical and pharmacological applications. Already in 1940, they were used to minimize osmotic shock after bleeding. Additional clinical uses include vaccine preparations and treatment of burn injuries, hemorrhagic shock, hypoproteinemia, and ascites resulting from liver cirrhosis(6, 7). HSA and BSA are also used in a range of biochemical procedures, such as immunological (e.g. ELISA), radioimmunological and immunoenzyme assays, as a blocking reagent, and as a molecular weight standard (4). Albumin is also widely used in molecular biology to stabilize reactants and prevent surface adhesion (8). Finally, HSA and BSA are used as supplements to cell-culture medium in the cultivated meat industry where recombinant albumin was shown to substitute the fetal bovine serum in the media of primary bovine satellite cells (9). Recent analyses suggest that this industry would require the production of millions of kilograms of recombinant albumin, exceeding the levels of production of many industrial enzymes(10). Clearly, economical and large-scale production of albumin may have important uses in a variety of basic and applied research settings.

Most albumin used in research and industry derives from animal or human plasma. Animal-sourced albumin, however, increases the risk of contamination from animal DNA and pathogens and suffers from batch-to-batch variability. This, and animal-welfare concerns, have encouraged the development of non-animal recombinant sources of albumin. Recombinant BSA and HSA from yeast and rice are used in some cases to replace plasma-derived albumin (11–13). Bacterial overexpression of proteins has significant advantages over yeast and plant-based expression in speed, costs, and yield. It would also enable screening albumin variants for desirable functional properties, such as enhanced ligand binding. Bacterial albumin expression is challenging, however, potentially due to its large size, multidomain organization, and 17 disulfide bonds. Indeed, bacterial expression of HSA was enabled only from inclusion bodies (14, 15), as a fusion to maltose-binding protein (16, 17), and with the use of chaperones (17). Expression in inclusion bodies and the use of expression tags may complicate downstream processing and reduce usefulness.

Here, we test whether albumins can be designed by the Protein Repair One Stop Shop (PROSS) algorithm (18) to improve their stability and enable bacterial expression while maintaining their functional properties. The PROSS stability design algorithm combines phylogenetic analysis of homologs and Rosetta atomistic design calculations to suggest a small set of mutants comprising dozens of stabilizing mutations (up to 20% of the protein). Previous studies in our lab and by others have shown orders of magnitude increases in bacterial expression and substantial improvement in heat tolerance even in designs of mammalian proteins (18–20). But we noted that improving the bacterial expressibility of disulfide-linked proteins is challenging(19). With 17 native disulfide bonds and several conformational states, serum albumin provides a challenging target for testing the ability of PROSS to improve the expressibility of dynamic disulfide-dependent proteins.

## Results

### Design strategy

Previous applications of PROSS often noted that designs with a greater number of mutations express at higher yields and are more heat tolerant(18, 19). We therefore decided to design several albumin variants with different mutational profiles for different uses. The HSA designs were based on PROSS design #7, and the BSA designs were based on design #6 of PROSS. In the most conservative design (HSA1 and BSA1), we allowed design only in the protein core and away from any binding sites. In the next design (HSA2 and BSA2), we also enabled design in the binding pockets, and in HSA3 and BSA3, we allowed mutations throughout the protein (**Table 1, Fig. 2, Supp. Tables 1 and 2**). We envision that Design 1 may find uses in settings that require the most conserved albumin, for instance, where binding sites must be completely conserved and no new binding sites may be introduced. Design 2 could be used when all solvent-accessible surfaces must remain intact, such as in raising albumin-targeting antibodies. Design 3 could be used in cases where albumin is used for its osmotic or non-specific surface-binding properties. Furthermore, crystallographic analyses reveal two major conformations for albumin (compact and myristate-bound), and in the design process, we selected only mutations that were observed in designs based on both structures to maintain the ability of the variants to change conformation.

**Table 1:** Number of mutations, stability, expression levels and binding affinities of HSA and BSA variants.

| Protein | # mutations<br>(% protein<br>mutated) | Melting<br>temperature<br>nanoDSF (DSC),<br>°C | Bacterial<br>expression,<br>mg/L culture | Binding affinity,<br>μM |
| --- | --- | --- | --- | --- |
| Wild type HSA<br>(commercial) | 0 | 60 | <2 | Warfarin:<br>19±6<br>Ketoprofen:<br>33±10 |
| HSA1<br>(no surface, no<br>binding site<br>mutations) | 16<br>(2.7) | 86 | 10-15 | Warfarin:<br>3.9±0.1<br>Ketoprofen:<br>7±2 |
| HSA2<br>(no surface<br>mutations) | 25<br>(4.3) | 90 | ~40 |  |
| HSA3 (mutations<br>all over the<br>protein) | 73<br>(12.5) | >95<br>(100.5) | ~100 |  |
| Wild type BSA<br>(commercial) | 0 | 58 | <2 |  |
| BSA1<br>(no surface, no<br>active site<br>mutations) | 16<br>(2.7) | 40-50 | <2 |  |
| BSA2<br>(no surface<br>mutations) | 29<br>(5.0) | 70 | ~20 |  |
| BSA3<br>(mutations all<br>over the protein) | 72<br>(12.3) | 85 | ~100 |  |

**Figure 2.**
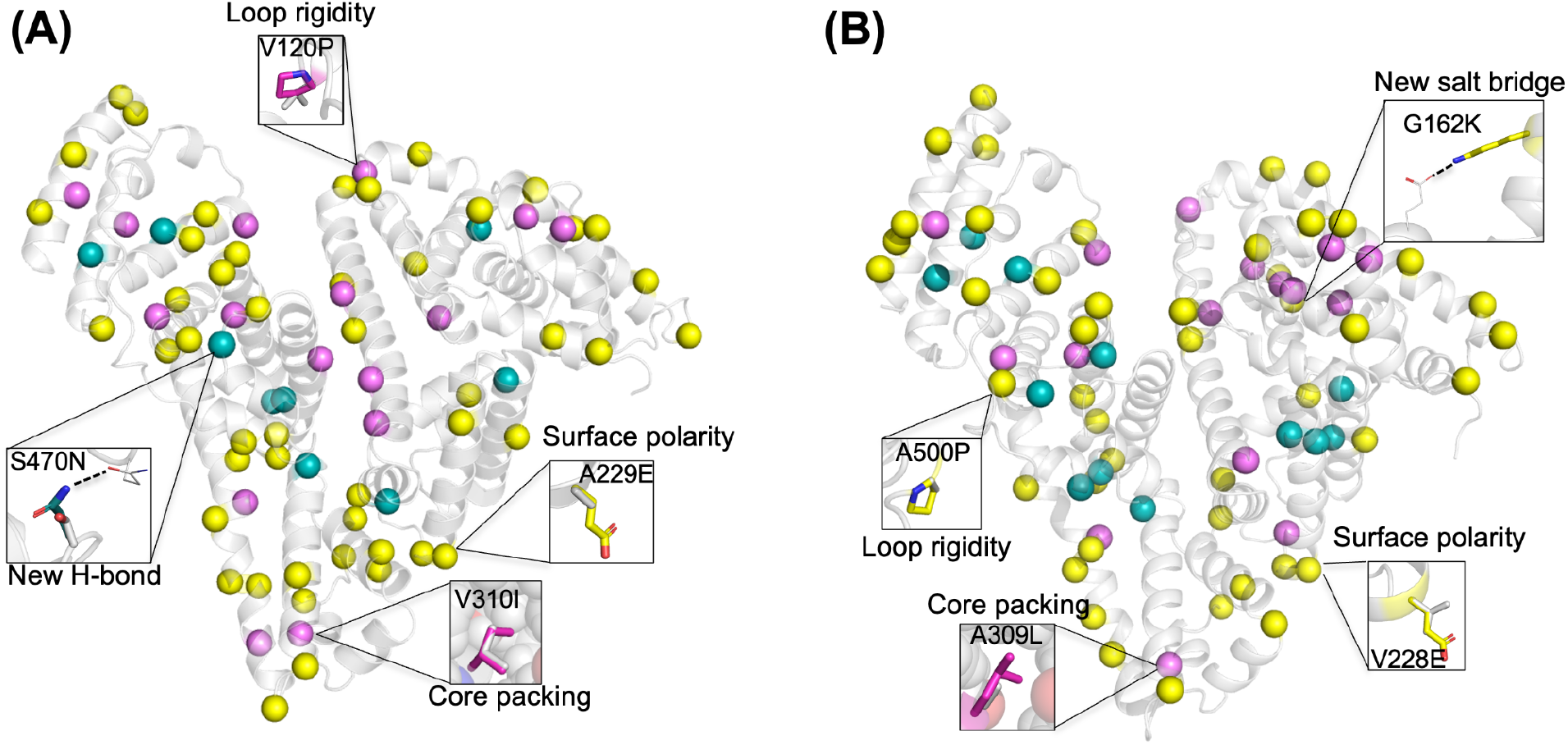
Mutations in stabilized human (**A**) and bovine (**B**) albumin designs. Design models with mutated positions indicated in spheres. Pink spheres - mutations in HSA1/BSA1; teal spheres - additional mutations in HSA2/BSA2, and yellow spheres - additional mutations in HSA3/BSA3. Thumbnails indicate possible stabilizing effects of selected mutations.

Since HSA and BSA are homologous, the PROSS mutational profiles partly overlapped (**Supp. Table 3**). Of the 119 mutations suggested by PROSS for HSA and BSA, 25 mutations (21%) were the same (same wild type identity in HSA and BSA mutated to the same identity). In 12 other cases (10%), the same mutation was designed both for HSA and BSA, although the wild type identities were different.

### Stabilized albumins for bacterial overexpression

As expected, wild type HSA did not express solubly in *E. coli*. We purified the HSA and BSA designs using Nickel-NTA affinity purification, finding that the HSA designs expressed robustly, and that as seen in a previous PROSS design study(19), expression levels increased with the number of stabilizing mutations (**Fig. 3A**) from ∼10 mg/L culture for HSA1 (16 mutations) to ∼100 mg/L culture for HSA3 (73 mutations). BSA trended similarly (**Fig. 3B**), but wild type BSA exhibited some soluble expression, and BSA1 showed only a modest increase in expression levels and was not characterized further.

**Figure 3.**
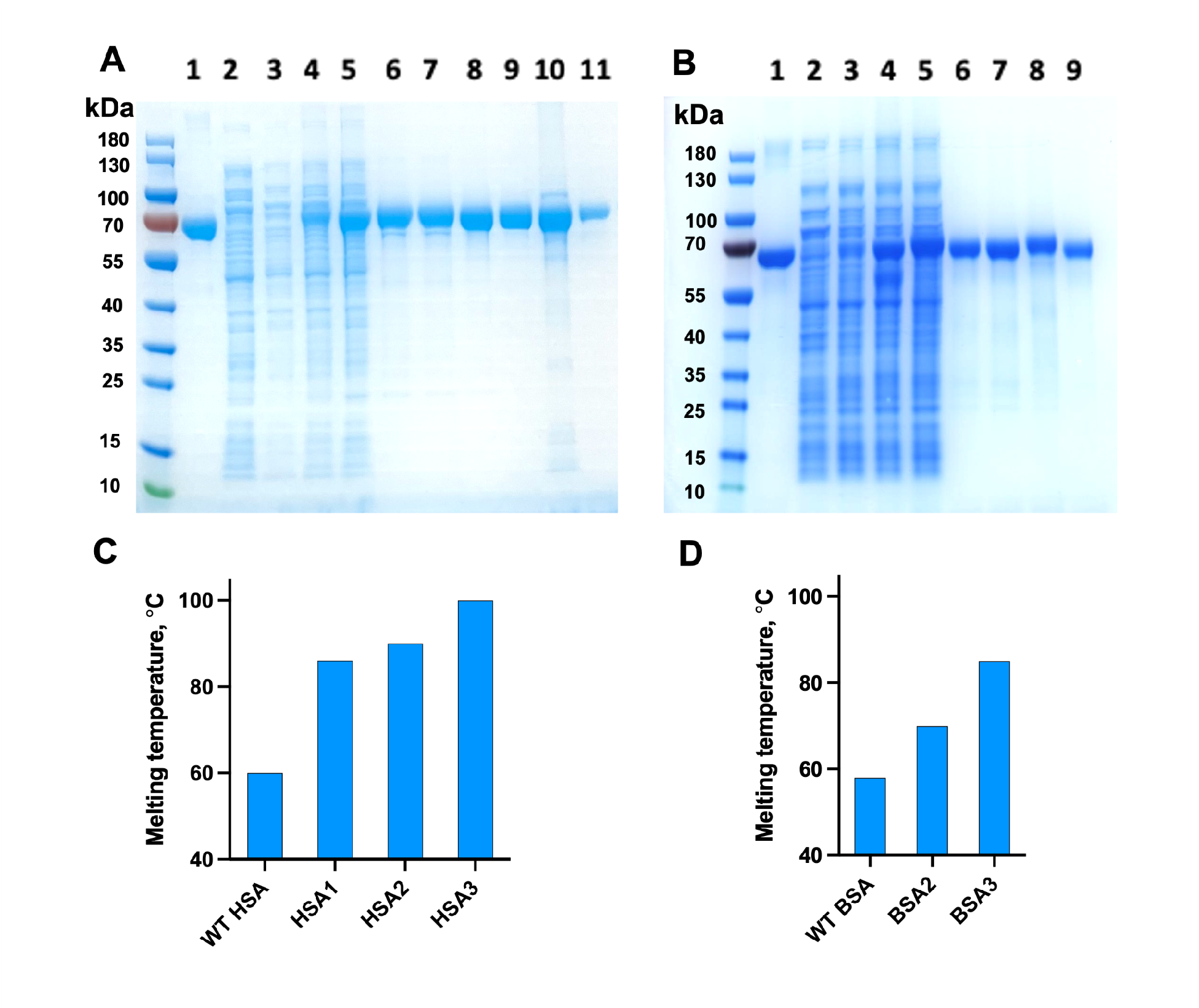
Soluble expression and thermal stability of designed HSA and BSA variants. **A**. SDS-PAGE gel of HSA variants. Commercial HSA (lane 1), supernatant fractions of His-tagged wild-type HSA (2), HSA1 (3), HSA2 (4), and HSA3 (5); purified His-tagged HSA1 (6), HSA2 (7), and HSA3 (8), purified tagless HSA1 from SUMO fusion (9), tagless HSA3 supernatant after heating for 5 minutes at 90°C (10), purified tagless HSA3 (11). **B**. SDS-PAGE gel of BSA variants. Commercial BSA (1), supernatant fractions of His-tagged wild-type BSA (2), BSA1 (3), BSA2 (4), and BSA3 (5); purified His-tagged BSA1 (6), BSA2 (7), and BSA3 (8), purified tagless BSA2 from SUMO fusion (9). **C**. Melting temperatures of HSA variants. **D**. Melting temperatures of BSA variants.

We next examined the apparent thermal stability of the designs compared to commercial plasma-derived HSA and BSA samples (**Table 1, Fig. 3C-D**). The apparent melting temperatures of the commercial samples were consistent with values reported earlier (approximately 60 °C) (21). As expected, thermal stability increased with the number of stabilizing mutations. Remarkably, the HSA designs exhibited apparent *T*_*m*_ values of 86-101°C, an increase of 26-40°C relative to the wild type HSA (**Fig. 3C, Supp. Fig. 1-5**). The BSA-based designs also improved apparent thermal stability by as much as 27°C relative to the wild type (**Fig. 3D, Supp. Fig. 6-9**).

Encouraged by the soluble expression and the increased thermostability of His-tagged HSA and BSA designs, we next designed tagless constructs of the least mutated variants. We expressed HSA1 and BSA2 fused to a C-terminal SUMO tag (22), which was cleaved scarlessly during purification, and applied size exclusion chromatography (**Fig. 3A-B**, lane 9). Because HSA3 resists boiling (>100°C), for this specific design, we substituted the affinity chromatography and cleavage steps with a heat-purification step. By heating the supernatant to 90°C and then cooling down to room temperature, we denatured and precipitated most protein impurities (**Fig. 3A**, lane 10). The remaining impurities were removed by size exclusion and ion exchange chromatography (**Fig. 3A**, lane 11), yielding over 100 mg protein from one liter of bacterial culture. Next, we analyzed the oligomeric state of the designs using size-exclusion chromatography. We found that the designs formed a mixture of monomeric and dimeric states, without larger aggregates (**Supp. Fig. 10-15**), similar to mammalian albumins (23). Tagless HSA1 and HSA3 were mostly monomeric, and tagless BSA2 had various proportions of monomeric and dimeric fractions (**Supp. Fig. 16-18**). Moreover, HSA and BSA designs remained mostly monomeric after four months of storage at 4°C (**Supp. Fig. 19**), and did not show any signs of aggregation.

Serum albumins carry a variety of ligands. We measured HSA1 binding to two site-specific drug ligands: warfarin, which binds to Sudlow site I, and ketoprofen, which binds to Sudlow site II (3). Despite 16 mutations from human albumin, HSA1 binding affinities for these ligands were similar and even better than those obtained for the commercial wild type HSA (**Table 1, Fig. 4**). The higher binding affinities observed in HSA1 compared to the wild type may be due to greater preorganization caused by protein stabilization or to uncontrollable differences in lipid composition and occupancy in the different albumin preparations.

**Figure 4.**
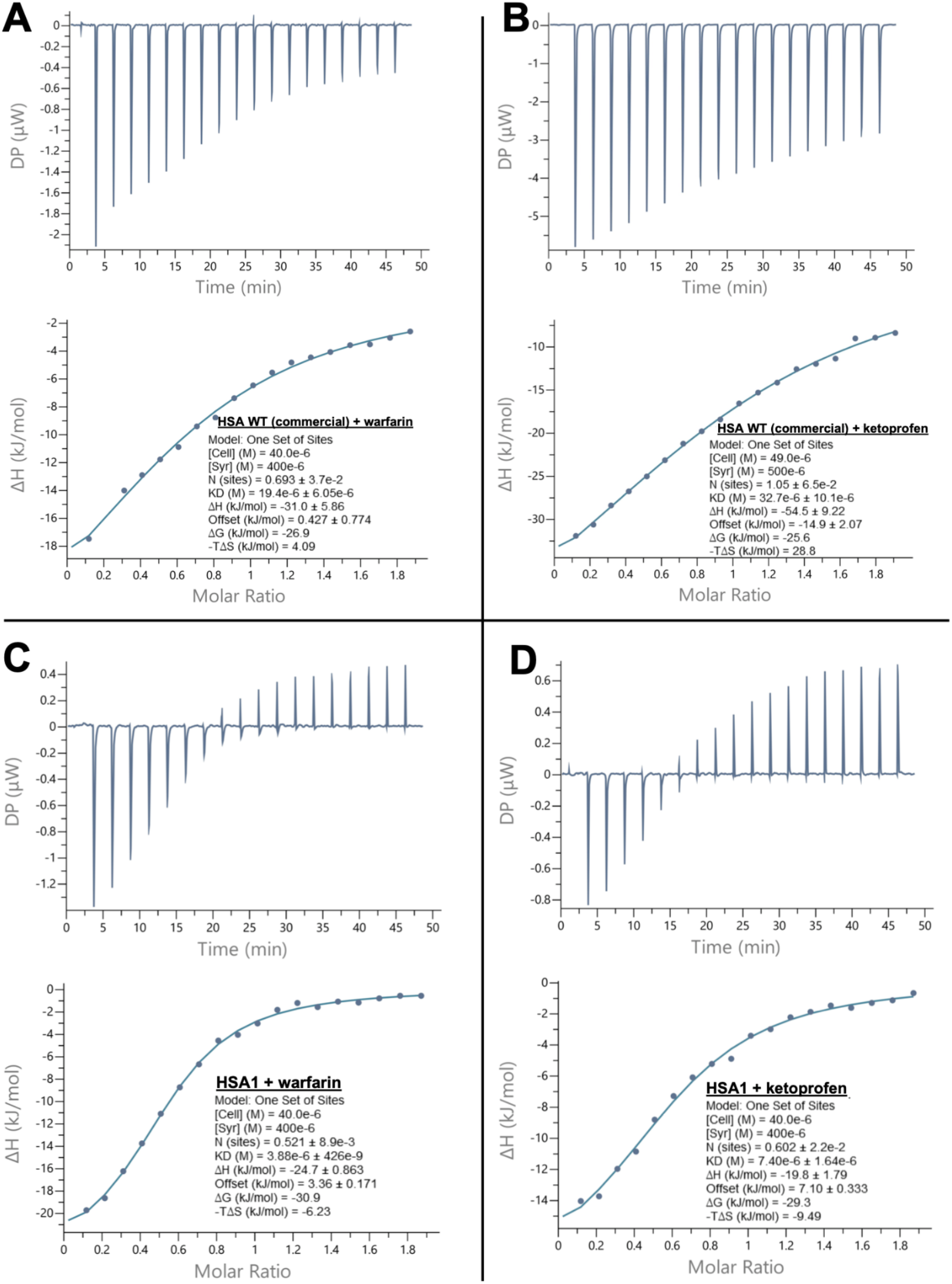
Ligand binding to human serum albumin variants, measured by ITC. (**A, C**) Warfarin binding to WT commercial HSA and HSA1, respectively. (**B, D**) Ketoprofen binding to commercial HSA and HSA1, respectively.

To verify the atomic accuracy of the design process, we determined the crystal structure of HSA1 in complex with warfarin. The structure revealed two monomers per asymmetric unit and diffracted to 2.0 Å resolution with warfarin occupying its native binding site (**Fig.5**), as in plasma-derived HSA (PDB entry 2bxd). Serum albumin is known to bind various fatty acids, among them myristate (24). Although myristate was not added to the crystallization reaction, four myristate molecules are present in the structure, closely matching the locations observed in the structure of wild type HSA (PDB entry 2bxi). The fatty-acid molecules we observe in our analysis may have derived from the bacterial expression host or from the growth medium. The structure of HSA1 is in excellent agreement (root mean square deviation 0.8Å) to the open conformation structure of plasma-derived wild type HSA with myristate (PDB entry 2bxi). Each of the 17 disulfide bonds is oxidized, and no significant changes in the side chain conformations are observed. These results demonstrate that stabilized HSA expressed in *E. coli* is properly folded, assumes a structure close to natively expressed HSA, and binds myristate in essentially the same mode as the wild type.

**Figure 5.**
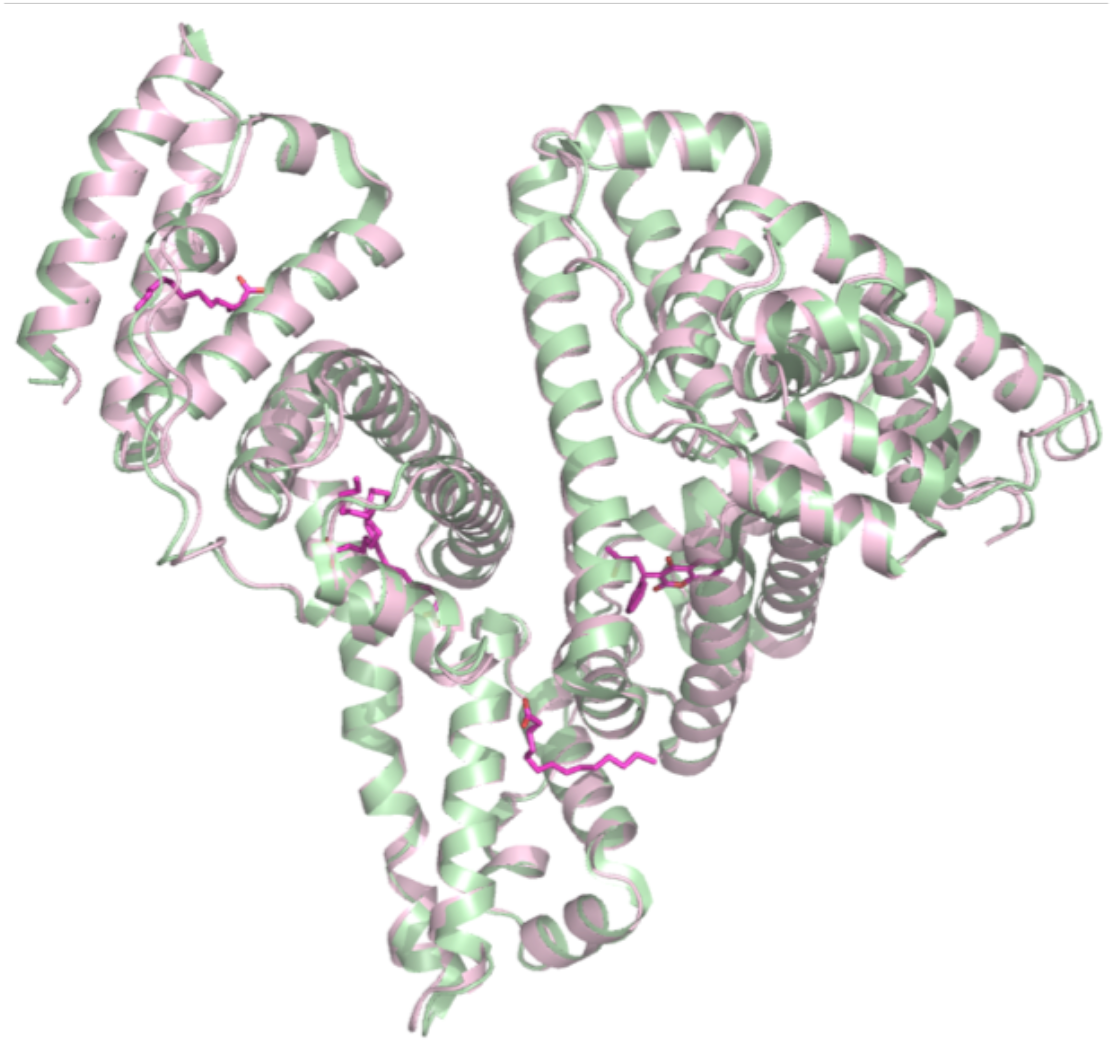
Crystal structure of HSA1 (PDB entry 8a9q; pink) overlaid on the structure of HSA with myristate (green, PDB entry 2bxi). The four myristate molecules and warfarin from the HSA1 structure are shown as magenta sticks.

### Stabilized HSA and BSA variants in cell culture and in vitro applications

To test whether the albumin designs are not toxic to human cells and can be used in cell culture medium, we added them to the growth medium of HEK293T cells and hybridoma cells and measured the number of cells and their viability. In the case of HEK293T cells, there was no adverse effect of HSA1, HSA3, or BSA2 variants, similar to commercial plasma-derived HSA and BSA (**Fig. 6A-B**). Similarly, the HSA and BSA designs did not have any adverse effect on the number of live hybridoma cells or on antibody-secretion levels (**Fig. 6C-D**).

**Figure 6.**
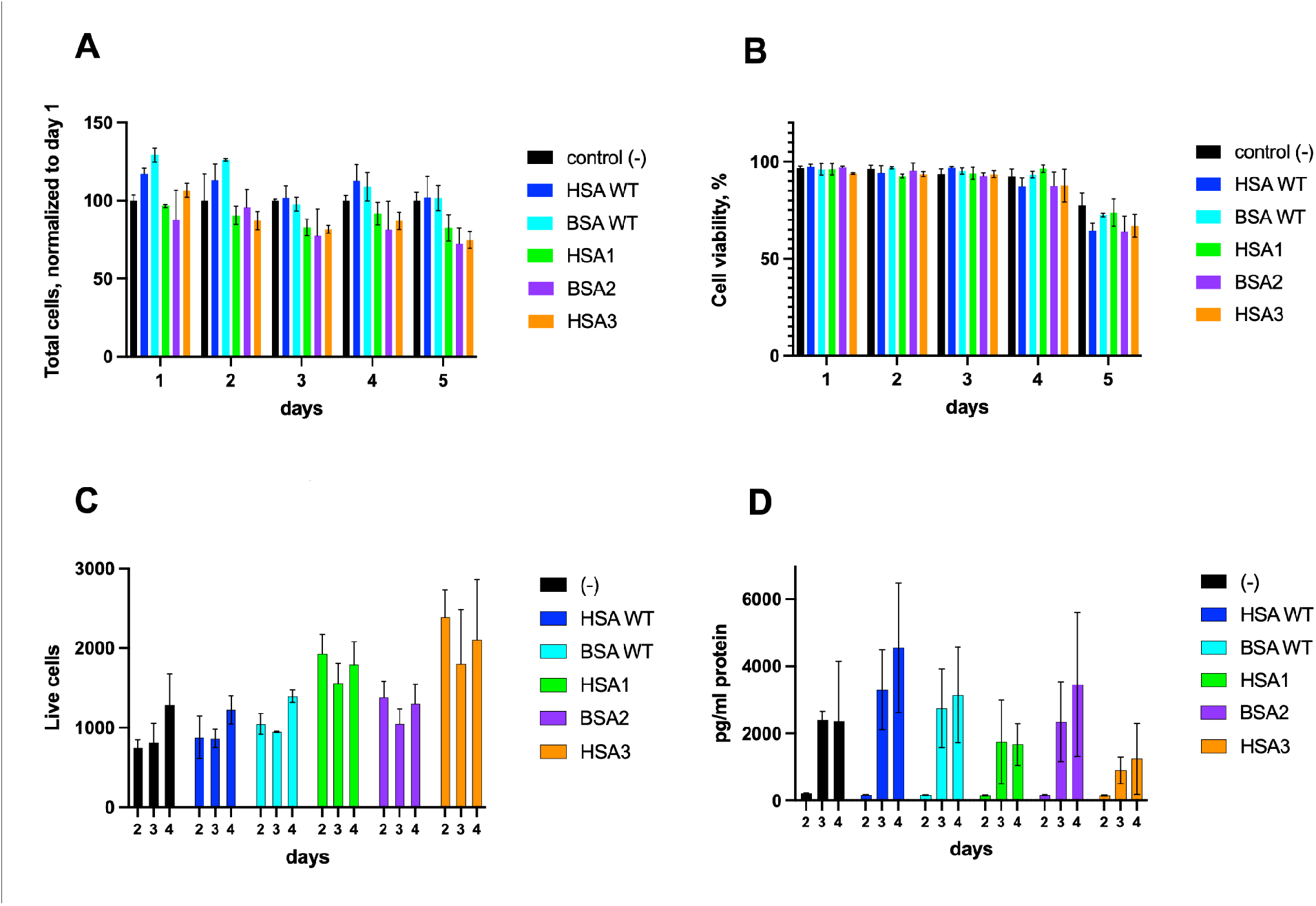
Stabilized albumin designs in cell culture medium. **(A)** Normalized number of HEK293T cells and **(B)** viability with 1 mg/ml of albumin designs (or plasma-sourced WT albumin) in medium with 5% FBS. **(C)** Live hybridoma cells and **(D)** antibody secretion with 1 mg/ml albumin designs (or animal-sourced albumin) in medium with 1% horse serum.

To test whether the designed albumins could be used in molecular biology applications, we performed restriction reactions and compared a buffer containing commercial BSA with a buffer supplemented with our stabilized albumins, HSA1, HSA3, or BSA2. In one case, pET28b(+) plasmid was digested with the restriction enzyme *BsiEI* (**Fig. 7**), and in another, pET29b(+) was digested with *NcoI* and *XhoI* (**Supp. Fig. 20**). In both cases, the designs and commercial BSA performed equally well, producing the same restriction pattern with the same efficiency. The albumin designs may be considered for use in reactions performed at extreme temperatures, such as PCR reactions, because their apparent melting temperatures are 70-100°C.

**Figure 7.**
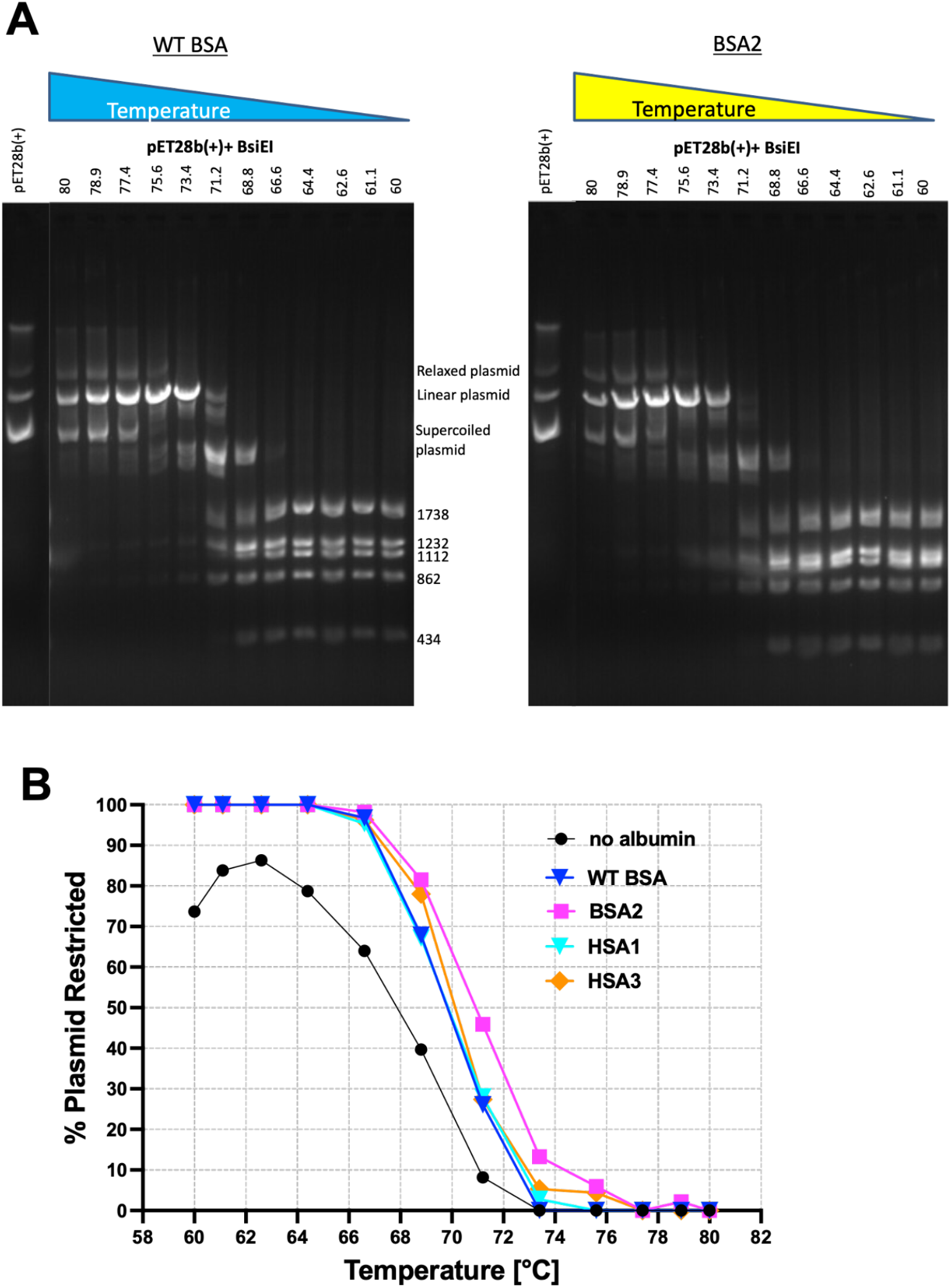
(**A)** Restriction of plasmid pET28b(+) with *BsiEI* in a buffer containing plasma-sourced WT BSA (left) or the BSA2 design (right). (**B)** % of restricted plasmid as a function of temperature in buffer containing albumin variants.

## Discussion

Human and bovine serum albumins have been widely used in a variety of clinical and biochemical applications for decades. Additionally, overexpression of recombinant albumin is likely to become a major bottleneck in the cultivated meat industry. In this study, we designed stabilized variants of HSA and BSA that are highly expressed in *E. coli* cells. Simple and fast expression and purification protocols make these variants amenable to cost-effective, large-scale production for research and industrial applications. The increased thermal stability may be useful in applications performed at high temperatures such as PCR or certain DNA restriction reactions. The ease of genetic manipulation and protein production in *E. coli* opens the way to design albumins with enhanced activities.

In recent years, the PROSS stability design algorithm has been successfully applied to a variety of enzymes and binding proteins(19, 25, 26). Designs typically exhibited higher melting temperatures and expression levels with minimal change in activity profile. The application of PROSS to serum albumins presented special challenges due to their large size, disulfide bonds, flexibility, and multiple ligand-binding sites. These complexities demanded an expert-driven analysis of the designed mutations to ensure that they are compatible with the two main conformational states and do not interfere with the binding pockets. Despite these challenges, we observed larger gains in thermal stability (exceeding the boiling point of water in one case) than in any previous study using PROSS. We speculate that the large number of disulfide bonds in serum albumins potentiates these proteins for very high stability but that due to the lack of evolutionary pressure, they do not exhibit such high stabilities in any natural homolog(27). According to this perspective, PROSS implemented mutations that enable the proteins to reach their high potential stability. Our study demonstrates that even challenging dynamic, multidomain, and disulfide-linked proteins can be substantially improved, requiring only a minor experimental effort.

## Methods

### Computational design

HSA and BSA were stabilized using the PROSS2 web server (28), using the default settings. HSA design was performed on two structures: PDB entry 2bx8 - with 2 azopropazone ligands, and PDB entry 2bxi - with 7 myristate ligands and 2 azopropazone ligands. Three designs with reduced sets of mutations were constructed, based on PROSS design 7.

BSA design was performed on three structures (with different ligands and thus different conformations): PDB entry 6qs9 (with ketoprofen), 4jk4 (with 2-hydroxy 3,5-diiodobenzoic acid), and 4or0 (with naproxen). As in the case of HSA, three designs were tested experimentally, all based on 4or0 design 6.

Designs were deposited in AddGene under accession number #82824.

### Protein expression

Several constructs of HSA and BSA were tested. Wild type HSA and BSA, and the designed genes were ordered from Twist Bioscience and cloned into a pET29b vector with C-terminal 6xHis tag (His-tag version). HSA1 and BSA2 variants were also cloned into a pET28-SUMO vector with cleavable N-terminal His-bdSUMO tag (SUMO version), and HSA3 variant was also cloned into a pET29b vector with deleted 6-His tag (no-His version).

All the variants were transformed into SHuffle T7 cells (NEB) and plated on LB plates with kanamycin (kana) and spectinomycin (spec). Ten milliliters of 2YT medium supplemented with 50ug/ml kana and 50ul/ml spec were inoculated with a single colony and grown overnight at 37°C. Then, 2YT medium supplemented with kana and spec (50-1500ml) was inoculated 1:100 with the overnight culture and grown at 37°C until OD_600_ of ∼0.6. Overexpression was induced with 0.3mM IPTG, the cultures were grown for 18-20h/20°C, harvested (5000rpm/10min/4°C) and the pellet was frozen at -20°C. The cells were dispersed in lysis buffer (PBS, Dulbecco’s phosphate buffered saline, Biological Industries #02-023-1A, pH 7.4) + 1:10000 benzonase (Sigma-Aldrich #E1014)) and lysed by sonication.

### Protein purification

Centrifuge Heraeus Multifuge X3 (Thermo Scientific) with rotor Fiberlite F15-8×50cy was used in protein purification procedures. His-tagged variants purification: The supernatant obtained after centrifugation (10,000rpm for 20 min at 4°C) was supplemented with 10mM imidazole and mixed with Ni-NTA resin (2-3h at 4°C), washed with 20mM imidazole in PBS and the protein was eluted with 250mM imidazole in PBS.

SUMO-tagged variants purification: The supernatant was purified on Ni-NTA resin as described before, but the elution was done by mixing overnight at 4°C in PBS + 1mM DTT with 5μg/ml bdSENP1 protease (22). The unbound fraction contained the tagless BSA/HSA variant.

Tagless HSA3 purification: The supernatant of HSA3 was incubated for 5 minutes at 90°C and cleared by centrifugation (10,000rpm/15min/4°C).

Variants expressed from cultures of 50-500ml were further purified by gel filtration on a Superdex 75 Increase 10/300 GL column (10 × 300 mm, Cytiva) using PBS + 150mM NaCl buffer. Variants purified in large-scale preps (>500ml culture) were purified by gel filtration using a HiLoad Superdex 200 pg column (26 × 600 mm, Cytiva) with PBS + 150mM NaCl buffer, and then by a HiPrep Q XL 16/10 anion-exchange column (Cytiva) using a gradient from 0 to 50% buffer B (20mM Tris HCl pH 8 + NaCl 1M) following buffer exchange to buffer A (20mM Tris HCl pH 8). Tagless proteins were similarly purified by gel filtration and anion exchange. Final protein purity was estimated to be >95% by SDS-PAGE.

### Thermal stability measurements

Thermal stability of BSA and HSA variants (His-tagged form) and of commercial wild type HSA and BSA (Sigma-Aldrich A1653 and A7638, respectively) was measured by nanoscale differential scanning fluorimetry (nanoDSF), in Prometheus NT.48 (NanoTemper). The temperature was ramped from 20°C to 95°C at 1°C/min. Thermal stability of HSA3 variant was also measured by differential scanning calorimetry (DSC) in VP-DSC instrument (Malvern), with heating from 20°C to 105°C at 1C°/min and cooling at the same rate.

### Ligand binding measurements

Ligand binding to commercial wild type HSA (non-defatted, Sigma-Aldrich A1653) and to HSA1 variant (His-tagged) was measured at 25°C by isothermal titration calorimetry (ITC, MicroCal ITC instrument, Malvern), in PBS supplemented with 1% DMSO. HSA concentration was maintained at 40μM in the cell, and 400μM of warfarin (Sigma-Aldrich A2250) or ketoprofen (Sigma-Aldrich K1751) were loaded into the injector. The proteins were titrated with 19 injections of 10ul ligands with a 3-min equilibration time spacing the injections. Titration of ligand to buffer (PBS with 1% DMSO) was used as a blank control, and one-site model was used to calculate the binding constants (KD).

### Crystallization and structure determination of HSA1 and Warfarin

Tagless HSA1 protein obtained from SUMO-HSA production was concentrated to 100mg/ml in Tris buffer pH 7.0 and supplemented with 1mM warfarin from 100mM stock in DMSO. Crystals of HSA1 and warfarin were obtained using the hanging-drop vapor diffusion method with a Mosquito robot (TTP LabTech). All datasets were collected under cryogenic conditions at the European Synchrotron Radiation Facility (ESRF), Grenoble, France at beamline ID30B. The crystals were grown from 0.05M NaCl, 10% PEG 4000 and 0.05 Tris pH 8.0. The crystals formed in the space group P1, with two monomers per asymmetric unit and diffracted to 2.0Å resolution. The integrated reflections were scaled using the AIMLESS program (29) from the CCP4i2 program suite (30). HSA1 structure determined by molecular replacement with PHASER (31) using the structure of human serum albumin in complex with aristolochic acid (PDB code 6HSC). All steps of atomic refinement were carried out with the CCP4i2/REFMAC5 program (32) and by PHENIX.refine (33). The models were built into 2mFobs - DFcalc, and mFobs - DFcalc maps by using the COOT program (34). The model was optimized using PDB_REDO (34, 35) and was evaluated with MOLPROBITY (36). Electron density revealed unambiguous density for the bound warfarin. Details of the refinement statistics of HSA1 and Warfarin structure are described in **Supp. Table 4**. The crystal structure was deposited in the PDB with PDB-ID code 8A9Q.

### Cell culture experiments

Tagless albumin variants were used for all the cell culture experiments. Hybridoma cells producing anti-GST monoclonal antibody (IgG1, Igk) were cultured for 2-4 days in the presence of 1% horse serum with or without 1mg/ml of the following proteins: commercial HSA, HSA1, HSA3, commercial BSA, and BSA2. The number of live cells and the levels of produced antibody were analyzed via flow cytometry and ELISA respectively. For flow cytometry, cells were collected every 24 hours stained with NucBlue™ Live Cell Stain (ThermoFisher LTD), according to manufacturer protocol and analyzed using LSRII cell analyzer plate reader. For ELISA, supernatants were collected every 24 hours and binding to GST was assessed using standard anti GST produced in our facility.

HEK293T adherent cells were grown in DMEM supplemented with GlutaMAX, NEAA and 5% FBS (all from Gibco) at 37°C, 5%CO_2_. To test the viability of the cultured cells in the presence of PROSS BSA2, HSA1 and HSA3 in comparison to the commercial BSA or HSA, respectively, cells (∼3×10^3^) were seeded in 24-well plate and followed each day for 5 days. All the albumin variants were added to the cells at 1mg/ml in duplicates, and their effect was analyzed. As a control, no protein was added to the growth medium. In each time point, cells were collected and the number of total cells, live cells and the fraction of viable cells were determined automatically using Brightfield Cell Counter DeNovix.

### DNA Restriction

pET29b vector with 900bp insert was restricted with *NcoI* and *XhoI* enzymes (NEB) overnight at 37°C, using the following buffers: rCutsmart buffer (NEB, contains 0.1mg/ml rBSA), buffer 4 (NEB, identical composition to rCutsmart buffer, but no BSA), and buffer 4 supplemented with HSA1, BSA2, and HSA3 variants (tagless, after gel filtration and ion exchange purification) at 0.1mg/ml.

Samples of pET28b(+) vector (1.5*μ*g) were restricted by *BsiEI* (10U, NEB) in 15*μ*l reaction volumes containing the Cutsmart buffer (NEB) or a reconstituted equivalent (50mM Potassium acetate, 20mM Tris-acetate, Mg-acetate, pH 7.9) supplemented with 0.1mg/ml of either commercial BSA (Sigma), HSA1, BSA2, or HSA3 variants (tagless, after gel filtration and ion exchange purification) at temperatures of 60-80°C for 15 minutes using a gradient PCR (SensoQuest). Samples were then supplemented with 5*μ*l DNA sample buffer (150mM TrisHCl pH 7.4, 5% SDS, 50% glycerol, 0.05% Bromophenol Blue), heated (60°C, 10 minutes), and resolved on a 2% agarose-TAE gel containing 0.5*μ*g/mL Ethidium Bromide for 1.5h at 120V.

### Analytical Gel Filtration Analysis

Samples of commercial BSA (Sigma) or tagless, purified variants BSA2, HSA1 or HSA3 (50*μ*l, 5.5mg) were loaded on an analytical size-exclusion HPLC column (BioSpec SEC-4000, Phenomenex) and eluted at 1ml/min using PBS buffer supplemented with 100mM NaCl, at RT.

## Abbreviations

HSA: human serum albumin
BSA: bovine serum albumin
PROSS: Protein Repair One Stop Shop
nanoDSF: nanoscale differential scanning fluorimetry
DSC: differential scanning calorimetry

## Accession numbers

The crystal structure of HSA1 has been deposited in the PDB with PDB ID **8A9Q**.

## Acknowledgements

We are grateful to Prof. Alessandro Angelini (Ca’ Foscari University of Venice, Italy) for helpful discussion and support. Research in the Fleishman lab was supported by the European Research Council through a Consolidator Award (815379), the Israel Science Foundation (1844), the Dr. Barry Sherman Institute for Medicinal Chemistry, and a donation in memory of Sam Switzer. We thank the Protein Analysis Unit for their help in experiments testing protein stability and ligand binding.

