## Supplementary Information for "Stable mammalian serum albumins designed for bacterial expression"

### Stable mammalian serum albumins for bacterial expression - Supplementary Information

**Supp. Table 1 - List of mutations in HSA variants**

**Supp. Table 2 - List of mutations in BSA variants**

**Supp. Table 3 - Comparison between HSA and BSA mutations**

**Supp. Table 4 - Refinement statistics details for HSA1 crystal structure**

**Supp. Fig. 1-4 - nanoDSF data of HSA variants**

**Supp. Fig. 5 - DSC of HSA3 variant**

**Supp. Fig. 6-9 - nanoDSF data of BSA variants**

**Supp. Fig. 10-19 - Gel filtration chromatograms of HSA and BSA variants**

**Supp. Fig. 20 - DNA restriction with NcoI-XhoI**

**Supp. Table 1.** List of mutations in HSA variants

| | Wild Type | HSA1-Conserved | HSA2-No surface mutations | HSA3 - raw | HSA_design_7 | Comments |
|---|---|---|---|---|---|---|
| # mutations | | 16 | 25 | 73 | 74 | |
| 33 | Q | | | K | K | |
| 38 | D | | | E | E | |
| 39 | H | L | L | L | L | |
| 42 | L | M | M | M | M | |
| 44 | N | | | K | K | |
| 52 | T | | | K | K | |
| 58 | S | | | T | T | |
| 76 | T | | | Q | Q | |
| 95 | E | | | D | D | |
| 99 | N | | | H | H | |
| 116 | V | | | E | E | |
| 120 | V | P | P | | | from PROSS run with 2bxi (with myristate) |
| 125 | T | | | K | K | |
| 136 | K | | N | N | N | partially exposed |
| 156 | F | Y | Y | Y | Y | new H-bond |
| 163 | A | | | K | K | |
| 172 | A | | | E | E | |
| 184 | E | | | A | A | |
| 187 | D | E | E | E | E | |
| 191 | A | | | E | E | |
| 198 | L | H | H | Y | Y | new H-bond |
| 202 | S | I | I | I | I | |
| 210 | A | | | | V | myristate binding site |
| 216 | V | | I | I | I | myristate binding site, but no clash with myristate expected |
| 227 | E | | | P | P | |
| 229 | A | | | E | E | |
| 231 | V | | | I | I | |
| 243 | T | | | K | K | |
| 244 | E | | | | A | |
| 250 | L | | | M | M | myristate binding site |
| 254 | A | | M | M | M | myristate binding site |
| 259 | D | | | K | K | |
| 286 | K | | | R | R | |
| 297 | E | | | D | D | |
| 298 | M | | | K | K | |
| 300 | A | | | E | E | |
| 304 | S | | | P | P | |
| 310 | V | I | I | | | mutation from designs 6 and 8, and PROSS run with 2bxi (with myristate) |
| 328 | G | | | A | A | myristate binding site |
| 329 | M | | | R | R | |
| 344 | V | | T | T | T | myristate binding site, no clash expected with myristate |
| 349 | L | | | I | I | |
| 355 | T | | | D | D | |
| 362 | A | | | K | K | |
| 364 | A | | | E | E | |
| 371 | A | S | S | | | mutation from PROSS run using talaris energy function, new H-bond |
| 374 | F | | | E | E | |
| 375 | D | | | E | E | |
| 379 | P | | | K | K | |
| 381 | V | I | I | | | from PROSS run with 2bxi (with myristate) |
| 384 | P | | | T | T | myristate binding site |
| 397 | Q | | | K | K | |
| 402 | K | | | Y | Y | myristate binding site |
| 409 | V | I | I | I | I | |
| 415 | V | | | M | M | myristate binding site |
| 419 | S | | | P | P | |
| 421 | P | | | D | D | |
| 426 | V | | | L | L | myristate binding site |
| 427 | S | A | A | T | T | |
| 440 | H | | L | L | L | partially exposed, in the cleft |
| 443 | A | | | E | E | |
| 446 | M | | | L | L | myristate binding site |
| 449 | A | | I | I | I | myristate binding site |
| 455 | V | I | I | I | I | |
| 470 | S | | N | N | N | Inside, not in a binding site |
| 486 | P | | | H | H | |
| 513 | I | | | L | L | Inside, not in a binding site, a bit close to myristate |
| 519 | K | E | E | E | E | |
| 524 | K | | | M | M | side chain not seen on 2bxi and 2bx8 structures |
| 527 | T | | | K | K | |
| 528 | A | | F | F | F | myristate binding site |
| 541 | K | | | E | E | |
| 547 | V | | I | | | myristate binding site, comes from design 9 and talaris designs |
| 552 | A | S | S | T | T | near MYR, no clash, T can make H-bond to carboxylic acid, new H-bond |
| 562 | D | | | E | E | |
| 564 | K | | | P | P | |
| 573 | K | | | S | S | |
| 576 | V | I | I | I | I | |
| 578 | A | | | K | K | |

Green - not surface, not binding sites mutations
yellow - myristate binding site mutations
Orange - surface mutations

**Supp. Table 2.** List of mutations in BSA variants

| | Wild Type | BSA1 - conserved | BSA2 - no surface mutations | BSA3 - raw | BSA_4or0_design_6 | Comment |
|---|---|---|---|---|---|---|
| # mutations | | 16 | 29 | 72 | | |
| 21 | G | A | A | **A** | | mutation from designs 8-9 |
| 28 | S | A | A | A | A | |
| 33 | Q | | | K | K | |
| 39 | H | **L** | **L** | **L** | **L** | |
| 42 | L | M | M | **M** | | mutation from designs 7-9 |
| 45 | E | | | D | D | |
| 52 | T | | | K | K | |
| 60 | A | | | P | P | |
| 63 | E | | | S | S | |
| 74 | L | | | | I | excluded, present only in design 9 in 6qs9 and 4jk4 design |
| 78 | A | | | E | E | |
| 79 | S | | | E | E | |
| 92 | E | | | S | S | |
| 109 | S | | | N | N | |
| 124 | D | | | K | K | |
| 128 | A | | | E | E | |
| 138 | L | I | I | I | I | |
| 158 | N | | | K | A | K is from design 8 |
| 162 | G | | | K | K | |
| 163 | V | I | I | **I** | | mutation from designs 8-9 |
| 174 | G | A | A | A | A | |
| 183 | T | | | A | A | |
| 184 | M | I | I | I | I | |
| 189 | L | | | K | K | |
| 202 | I | L | L | L | L | |
| 214 | S | | | | F | excluded, active site for diiodosalicylic acid in 4jk4 |
| 226 | E | | | P | P | |
| 228 | V | | | E | E | |
| 230 | V | I | I | I | I | |
| 240 | V | | I | I | I | active site for diiodosalicylic acid in 4jk4 |
| 241 | H | | Y | Y | Y | active site for diiodosalicylic acid in 4jk4 |
| 253 | A | | M | M | M | active site for myristate |
| 258 | D | | | K | K | |
| 260 | A | | V | V | V | active site for diiodosalicylic acid in 4jk4 |
| 285 | K | | | R | R | |
| 292 | V | | | | L | excluded, not present in designs based on 4jk4 and 6qs9 |
| 294 | K | | | N | F | N is from designs 7-9 |
| 296 | A | | | D | D | |
| 300 | N | | | D | D | |
| 305 | T | | | | V | excluded, the residue is exposed |
| 309 | A | L | L | L | L | |
| 321 | A | | | D | D | |
| 328 | S | | | R | R | |
| 344 | V | | L | L | L | active site for myristate |
| 351 | E | | | | V | excluded, exposed polar residue |
| 361 | A | | | K | K | |
| 371 | T | | | R | R | |
| 374 | D | | | E | E | |
| 378 | H | | | K | K | |
| 379 | L | | | H | H | |
| 380 | V | I | I | I | I | |
| 385 | N | | | E | | mutation from designs 7-9 and designs based on 4jk4 and |
| 387 | I | | | | V | excluded, not present in designs based on 4jk4 and 6qs9 |
| 408 | V | I | I | I | I | |
| 412 | R | | | K | K | |
| 414 | V | | M | M | M | active site for myristate |
| 418 | S | | | P | P | |
| 420 | P | | | D | D | |
| 425 | V | | I | I | | mutation from designs based on 6qs9 and 4jk4 |
| 426 | S | T | T | **T** | | mutation from designs 8-9 |
| 438 | T | | | Q | Q | |
| 439 | K | | L | L | L | partially exposed, in a cleft |
| 442 | S | | | E | E | |
| 443 | E | | | K | K | |
| 445 | M | | L | L | L | close to myristate in human SA structure 2bxi |
| 485 | P | | H | H | H | close to myristate in human SA structure 2bxi |
| 500 | A | | | P | P | |
| 503 | E | | | P | P | |
| 517 | D | | | P | P | |
| 518 | T | E | E | E | E | |
| 520 | K | | | | L | excluded because K had H-bond |
| 526 | T | | | K | K | |
| 527 | A | | F | F | F | active site for myristate |
| 546 | V | | I | I | | active site for myristate, mutation from design 9 and 6qs9 |
| 551 | V | | T | T | T | close to myristate in human SA structure 2bxi |
| 553 | F | | | | M | excluded, not present in designs based on 4jk4 and 6qs9 |
| 559 | A | | | K | K | |
| 561 | D | | | E | E | |
| 569 | V | | | E | E | |
| 575 | V | I | I | I | I | |
| 576 | V | | | E | E | |
| 577 | S | | | K | K | |
| 580 | T | | | A | A | |

Supp. Table 3. Comparison between HSA and BSA mutations

| | Wild Type HSA identity | HSA mutation | | Wild Type BSA identity | BSA mutation | | |
|---|---|---|---|---|---|---|---|
| 21 | A | | 21 | G | | | HSA already has Ala |
| 28 | A | | 28 | S | A | | HSA already has Ala |
| 33 | Q | K | 33 | Q | K | | same position, same mutation |
| 38 | D | E | 38 | | | | BSA already has Glu |
| 39 | H | L | 39 | H | L | | same position, same mutation |
| 42 | L | M | 42 | L | | | same position, same mutation |
| 44 | N | K | 44 | | | | adjacent position was mutated |
| 45 | E | | 45 | E | D | | adjacent position was mutated |
| 52 | T | K | 52 | T | K | | same position, same mutation |
| 58 | S | T | 58 | S | | | no mutation was suggested for BSA |
| 60 | E | | 60 | A | P | | no mutation was suggested for HSA |
| 63 | D | | 63 | E | S | | no mutation was suggested for HSA |
| 74 | L | | 74 | L | I | | no mutation was suggested for HSA |
| 76 | T | Q | 76 | K | | | no mutation was suggested for BSA |
| 78 | A | | 78 | A | E | | different conformation |
| 79 | T | | 79 | S | E | | different conformation |
| 92 | A | | 92 | E | S | | no mutation was suggested for HSA |
| 95 | E | D | 95 | | | | no mutation was suggested for BSA |
| 99 | N | H | 99 | | | | no mutation was suggested for BSA |
| 109 | N | | 109 | S | N | | HSA already has Asn |
| 116 | V | E | 116 | K | | | no mutation was suggested for BSA |
| 120 | V | P | 119 | P | | | BSA already has Pro |
| 125 | T | K | 124 | D | K | | same mutation, different initial identity |
| 129 | D | | 128 | A | E | | HSA already has Asp |
| 136 | K | N | 135 | G | | | no mutation was suggested for BSA |
| 139 | L | | 138 | L | I | | no mutation was suggested for HSA |
| 156 | F | Y | 155 | Y | | | BSA already has Tyr |
| 159 | K | | 158 | N | A | | no mutation was suggested for HSA |
| 163 | A | K | 162 | G | K | | same mutation, different initial identity |
| 164 | A | | 163 | V | I | | no mutation was suggested for HSA |
| 172 | A | E | 174 | G | A | | adjacent position was mutated |
| 184 | E | A | 183 | T | A | | same mutation, different initial identity |
| 185 | L | | 184 | M | I | | no mutation was suggested for HSA |
| 187 | D | E | 186 | E | | | BSA already has Glu |
| 190 | K | | 189 | L | K | | HSA already has K |
| 191 | A | E | 190 | T | | | no mutation was suggested for BSA |
| 198 | L | Y | 197 | L | | | no mutation was suggested for BSA |
| 202 | S | I | 201 | S | | | no mutation was suggested for BSA |
| 203 | L | | 202 | I | L | | HSA already has Leu |
| 210 | A | V | 209 | A | | | no mutation was suggested for BSA |
| 215 | A | | 214 | S | F | | different conformation |
| 216 | V | I | 215 | | | | different conformation |
| 227 | E | P | 226 | E | P | | same position, same mutation |
| 229 | A | E | 228 | V | E | | same mutation, different initial identity |
| 231 | V | I | 230 | V | I | | same position, same mutation |
| 241 | V | | 240 | V | I | | no mutation was suggested for HSA |
| 242 | H | | 241 | H | Y | | no mutation was suggested for HSA |
| 243 | T | K | 242 | K | | | BSA already has Lys |
| 244 | E | A | 243 | E | | | no mutation was suggested for BSA |
| 250 | L | M | 249 | L | | | no mutation was suggested for BSA |
| 254 | A | M | 253 | A | M | | same position, same mutation |
| 259 | D | K | 258 | D | K | | same position, same mutation |
| 261 | | | 260 | A | V | | no mutation was suggested for HSA |
| 286 | K | R | 285 | K | R | | same position, same mutation |
| 293 | V | | 292 | V | L | | different conformation |
| 295 | N | | 294 | K | F | | HSA already has Asn |
| 297 | E | D | 296 | A | D | | same mutation, different initial identity |
| 298 | M | K | 297 | I | | | different conformation |
| 300 | A | E | 299 | E | | | BSA already has Glu |
| 301 | D | | 300 | N | D | | HSA already has Asp |
| 304 | S | P | 303 | P | | | BSA already has Pro |
| 306 | A | | 305 | T | V | | no mutation was suggested for HSA |
| 310 | V | I | 309 | A | L | | same position, similar mutation |
| 322 | A | | 321 | A | D | | different conformation |
| 328 | G | A | 327 | G | | | different conformation |
| 329 | M | R | 328 | S | R | | same mutation, different initial identity |
| 344 | V | T | 343 | S | | | no mutation was suggested for HSA |
| 345 | L | | 344 | V | L | | HSA already has Leu |
| 349 | L | I | 348 | L | | | different conformation |

| | | | | | | | |
|---|---|---|---|---|---|---|---|
| 355 | T | D | 354 | A | | | no mutation was suggested for HSA |
| 362 | A | K | 361 | A | K | | same position, same mutation |
| 364 | A | E | 363 | D | | | different conformation |
| 372 | K | | 371 | T | R | | different conformation |
| 374 | F | E | 373 | F | | | different conformation |
| 375 | D | E | 374 | D | E | | same position, same mutation |
| 379 | P | K | 378 | H | K | | same mutation, different initial identity |
| 380 | L | | 379 | L | H | | different conformation |
| 381 | V | I | 380 | V | I | | same position, same mutation |
| 384 | P | T | 383 | P | | | different conformation |
| 386 | N | | 385 | N | | | different conformation |
| 388 | I | | 387 | I | V | | different conformation |
| 397 | Q | K | 396 | K | | | BSA already has Lys |
| 402 | K | Y | 401 | G | | | different conformation |
| 409 | V | I | 408 | V | I | | same position, same mutation |
| 413 | K | | 412 | R | K | | HSA already has Lys |
| 415 | V | M | 414 | V | M | | same position, same mutation |
| 419 | S | P | 418 | S | P | | same position, same mutation |
| 421 | P | D | 420 | P | D | | same position, same mutation |
| 426 | V | L | 425 | V | | | same position, similar mutation |
| 427 | S | T | 426 | S | | | same position, same mutation |
| 439 | K | | 438 | T | Q | | different conformation |
| 440 | H | L | 439 | K | L | | same mutation, different initial identity |
| 443 | A | E | 442 | S | E | | same mutation, different initial identity |
| 444 | K | | 443 | E | K | | different conformation |
| 446 | M | L | 445 | M | L | | same position, same mutation |
| 449 | A | I | 448 | T | | | no mutation was suggested for BSA |
| 455 | V | I | 454 | L | | | no mutation was suggested for BSA |
| 470 | S | N | 469 | S | | | no mutation was suggested for BSA |
| 486 | P | H | 485 | P | H | | same position, same mutation |
| 501 | E | | 500 | A | P | | different conformation |
| 504 | A | | 503 | E | P | | different conformation |
| 513 | I | L | 512 | I | | | no mutation was suggested for BSA |
| 518 | E | | 517 | D | P | | different conformation |
| 519 | K | E | 518 | T | E | | same mutation, different initial identity |
| 524 | K | M | 523 | K | | | different conformation |
| 527 | T | K | 526 | T | K | | same position, same mutation |
| 528 | A | F | 527 | A | F | | same position, same mutation |
| 541 | K | E | 540 | E | | | BSA already has Glu |
| 547 | V | I | 546 | V | | | same position, same mutation |
| 552 | A | T | 551 | V | T | | same mutation, different initial identity |
| 560 | K | | 559 | A | K | | HSA already has Lys |
| 562 | D | E | 561 | D | E | | same position, same mutation |
| 564 | K | P | 563 | D | | | different conformation |
| 570 | E | | 569 | V | E | | HSA already has Glu |
| 573 | K | S | 572 | P | | | no mutation was suggested for BSA |
| 576 | V | I | 575 | V | I | | same position, same mutation |
| 577 | A | | 576 | V | E | | Glu was suggested in some variants |
| 578 | A | K | 577 | S | K | | same mutation, different initial identity |
| 581 | A | | 580 | T | A | | HSA already has Ala |

**Supp. Table 4.** Data collection and refinement statistics for HSA1

| | |
|---|---|
| **Data Collection** | |
| PDB code | 8a9q |
| Space group | *P1* |
| Cell dimensions: | |
| a,b,c (Å) | 38.26 92.13 95.55 |
| α,β,γ (˚) | 74.3, 89.3, 80.0 |
| No. of copies in a.u. | 2 |
| Resolution (Å) | 46.02-2.00 |
| Upper resolution shell (Å) | 2.04-2.00 |
| Unique reflections | 74,409 (4,259) |
| Completeness (%)<br>Multiplicity | 89.1 (83.4)<br>10.2 (10.2) |
| Average I/σ(I) | 5.7 (1.1) |
| R-pim<br>CC1/2 | 0.05816 (0.2589)<br>0.945 (0.86) |
| **Refinement** | |
| Resolution range (Å) | 46.02-2.00 |
| No. of reflections | 70,736 |
| No. of reflections in test set | 3,634 |

| | |
|---|---|
| R-working / R-free | 0.2258 / 0.2684 |
| No. of protein atoms | 8923 |
| No. of water molecules | 12 |
| Overall average B factor ($Å^2$) | 42.98 |
| Root mean square deviations: | |
| - bond length (Å) | 0.017 |
| - bond angle (°) | 2.17 |
| **Ramachandran Plot** | |
| Most favored (%) | 96.55 |
| Additionally allowed (%) | 3.36 |
| Disallowed (%) | 0.09 |

* Values in parentheses refer to the data of the corresponding upper resolution shell

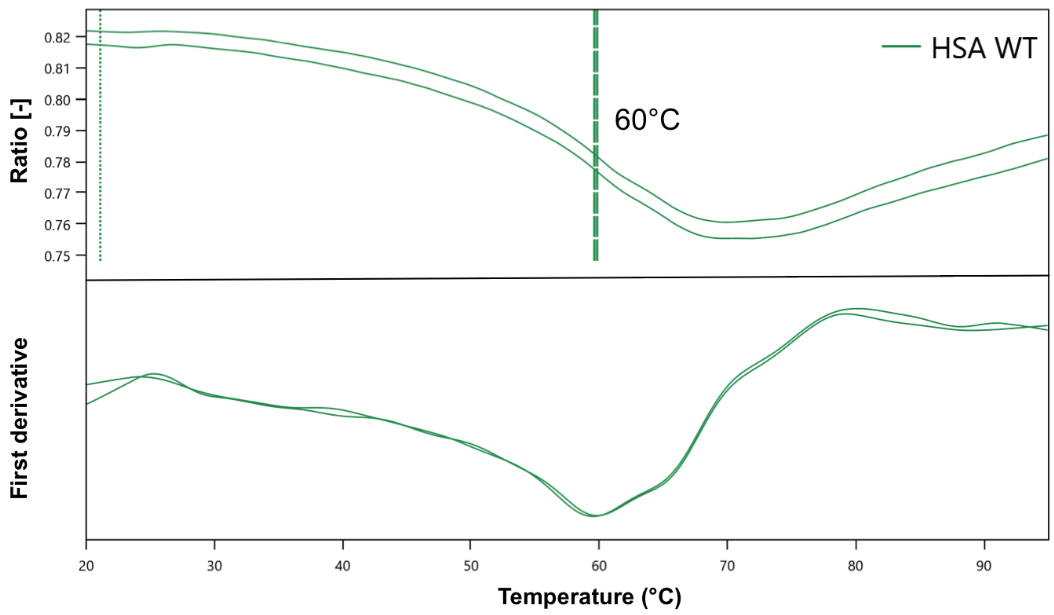


**Supp. Fig. 1.** nanoDSF of HSA WT variant

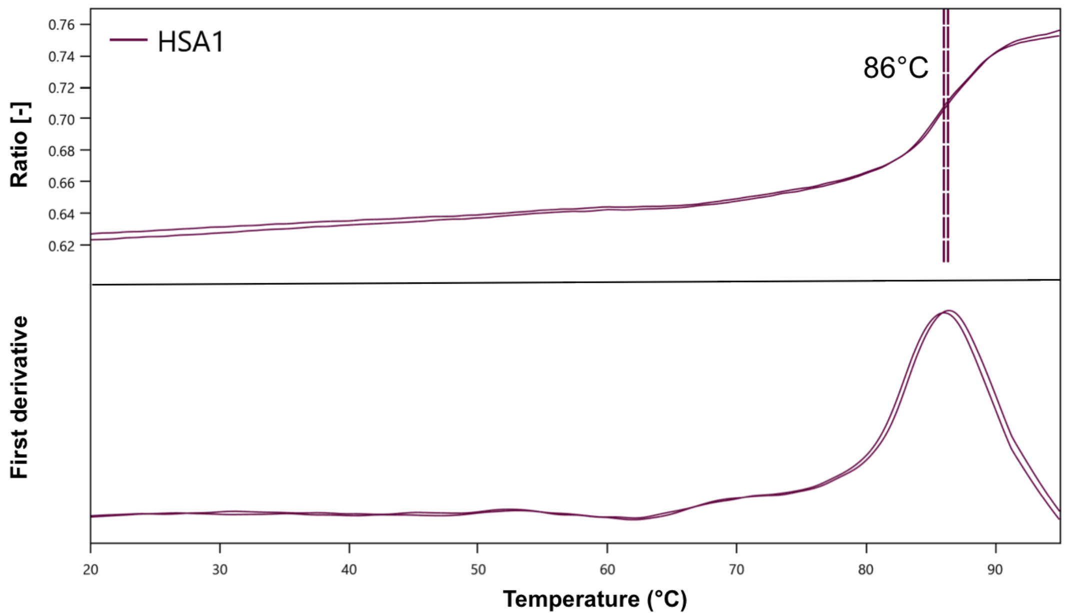


**Supp. Fig. 2.** nanoDSF of HSA1 variant

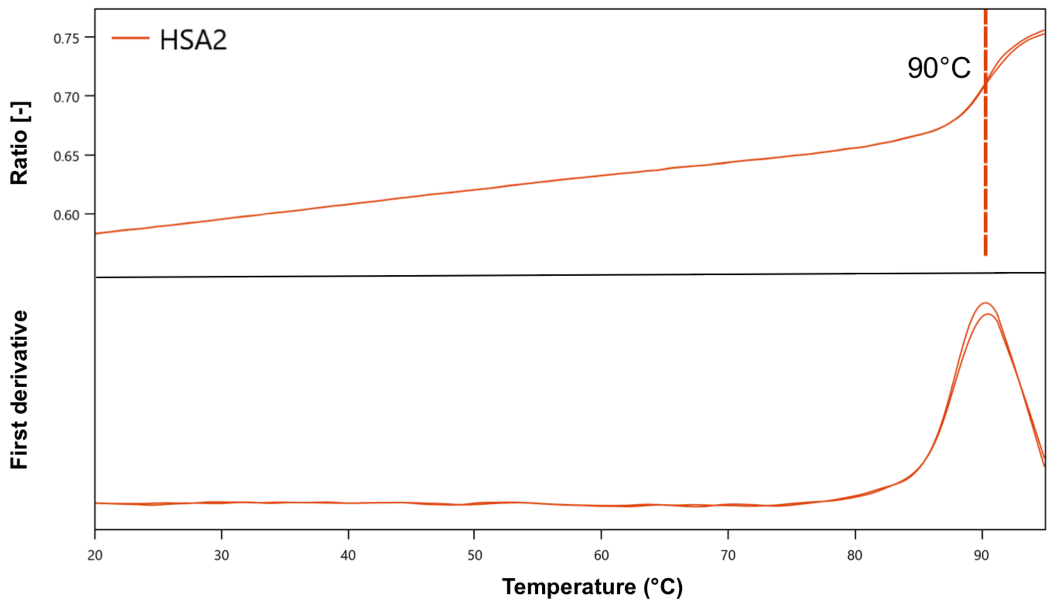


**Supp. Fig. 3.** nanoDSF of HSA2 variant

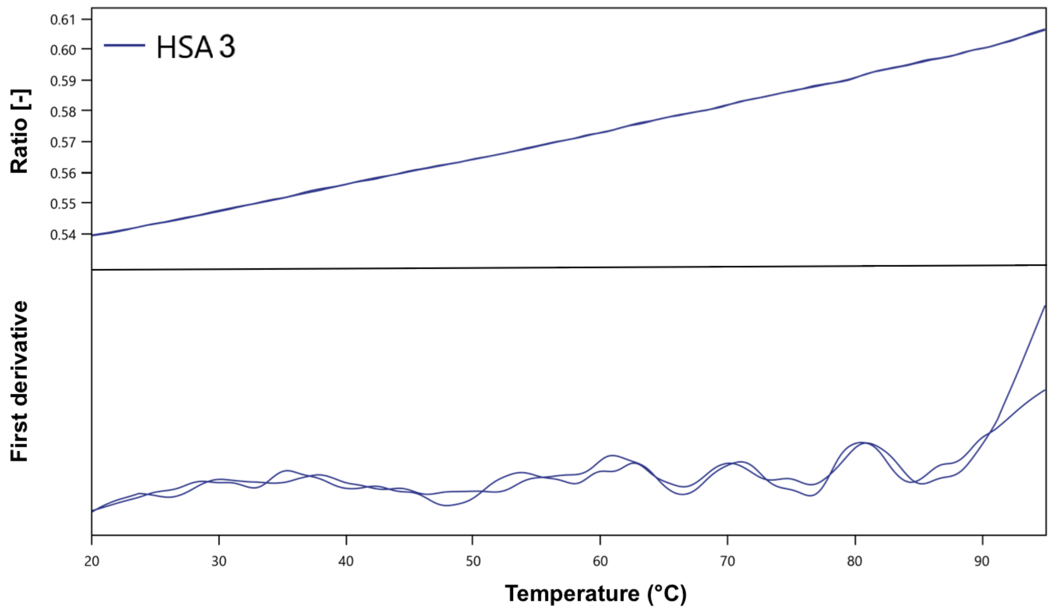


**Supp. Fig. 4.** nanoDSF of HSA3 variant you have a better thermogram that reaches 105 C

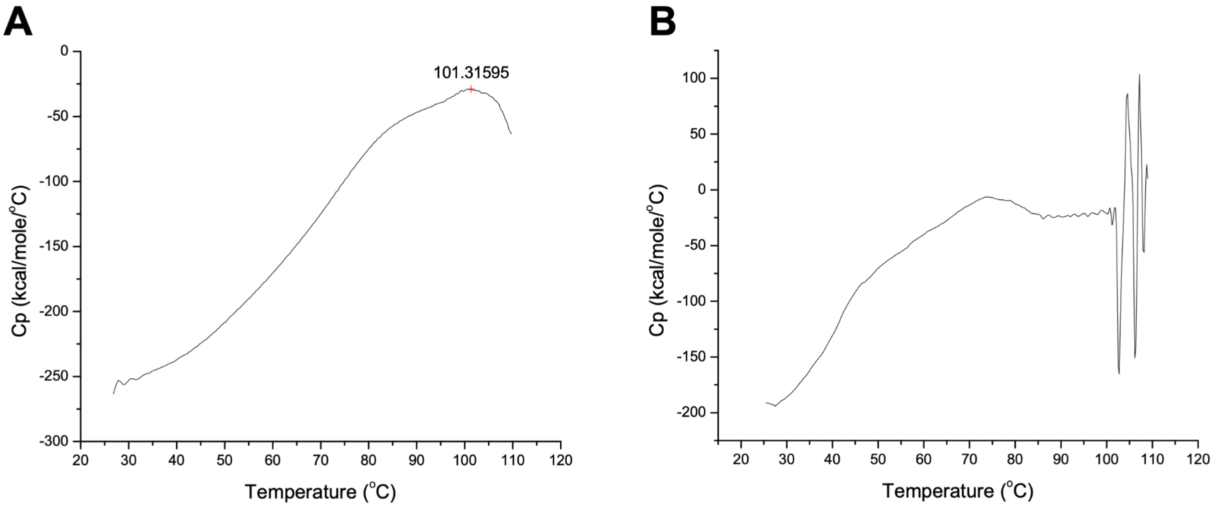


**Supp. Fig. 5.** DSC of HSA3 variant. **A.** Heating (unfolding). **B.** Cooling (refolding).

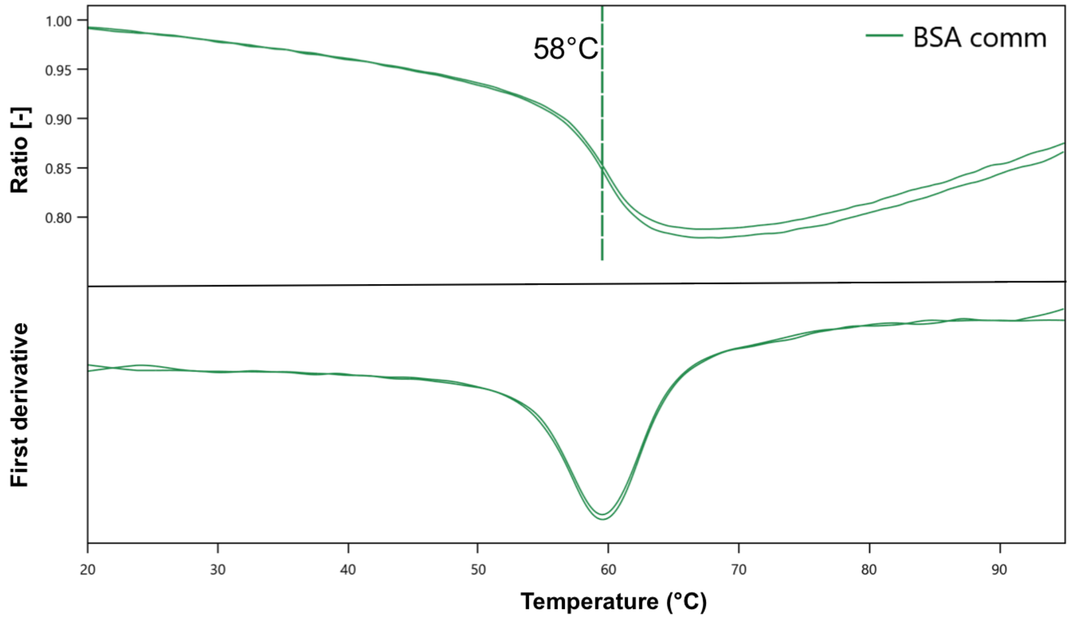


**Supp. Fig. 6.** nanoDSF of BSA WT variant (commercial BSA)

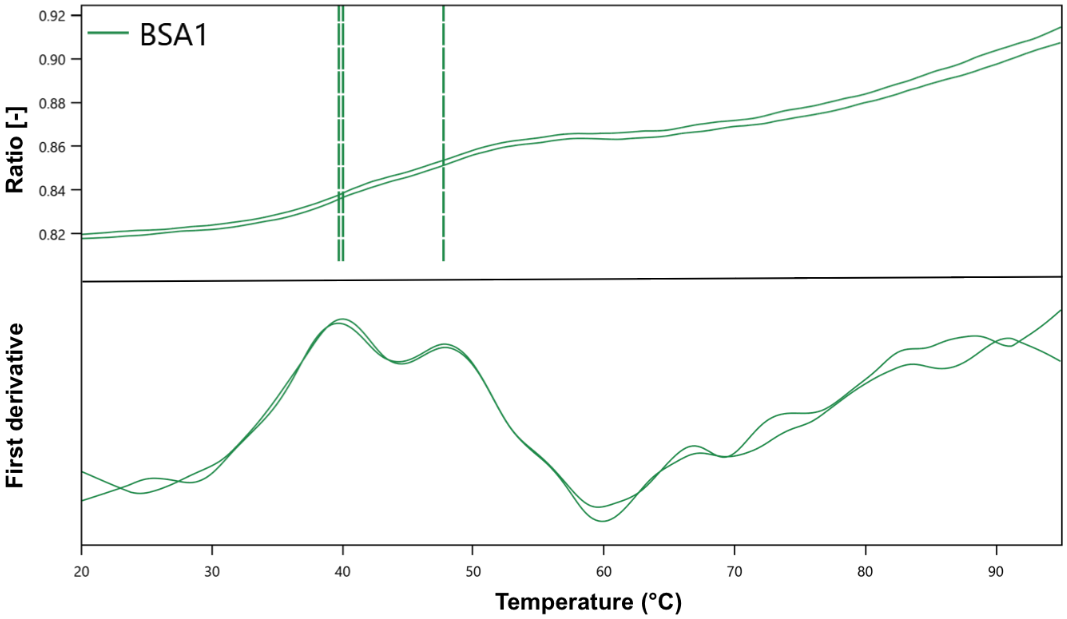


**Supp. Fig. 7.** nanoDSF of BSA1 variant (no melting observed, probably misfolded)

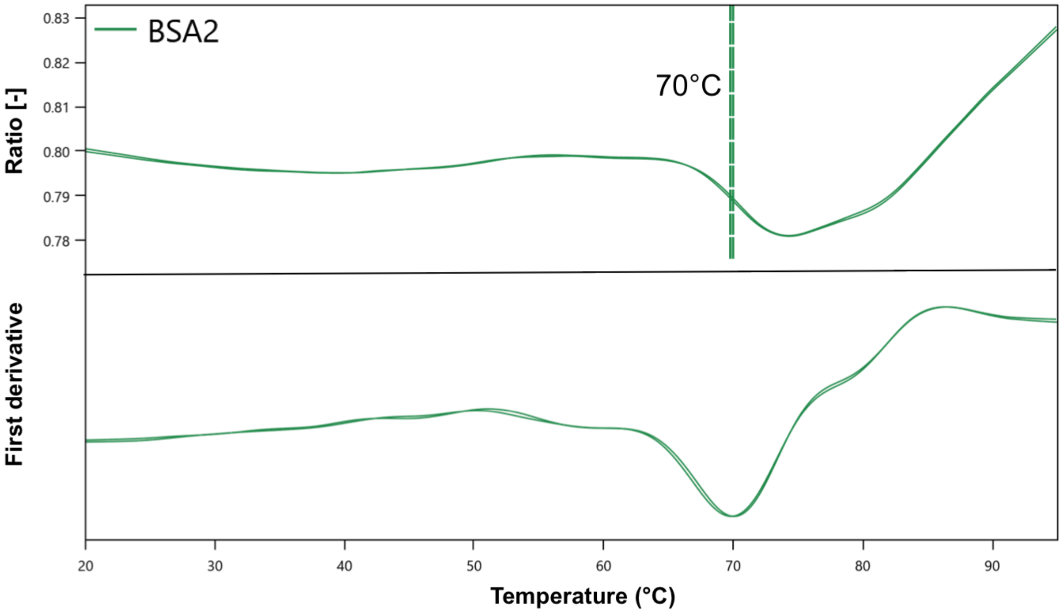


**Supp. Fig. 8.** nanoDSF of BSA2 variant

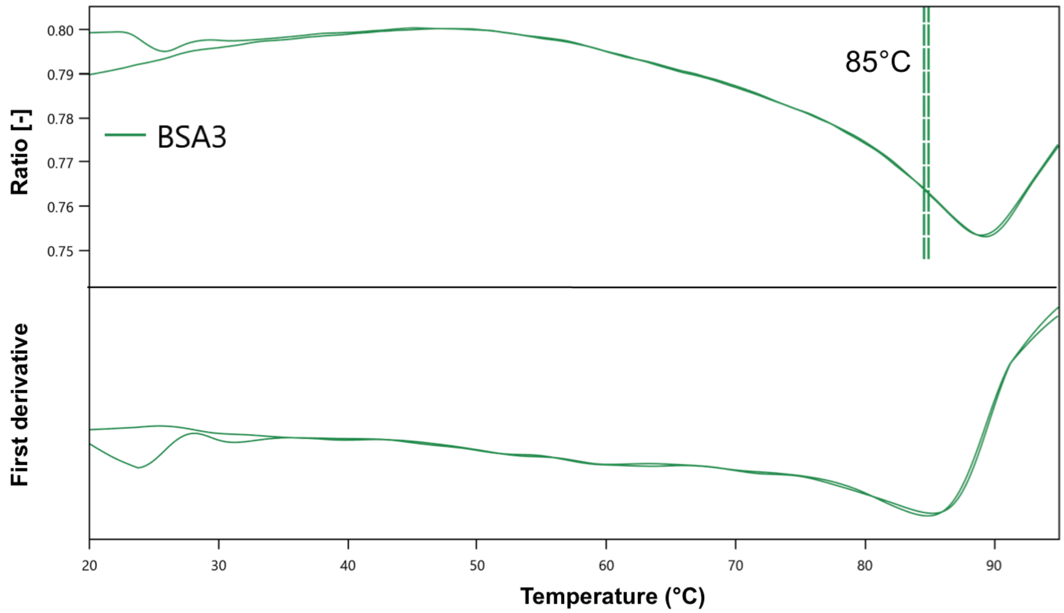


**Supp. Fig. 9.** nanoDSF of BSA3 variant

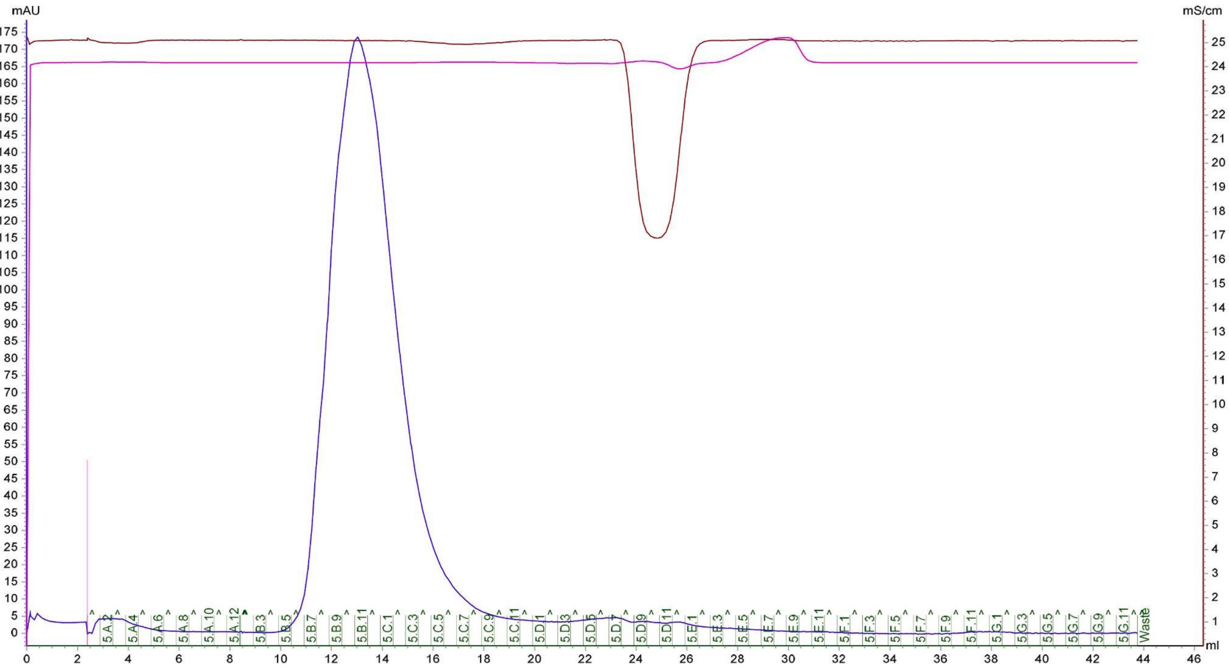


**Supp. Fig. 10.** Gel filtration chromatogram of His-tagged HSA1 variant

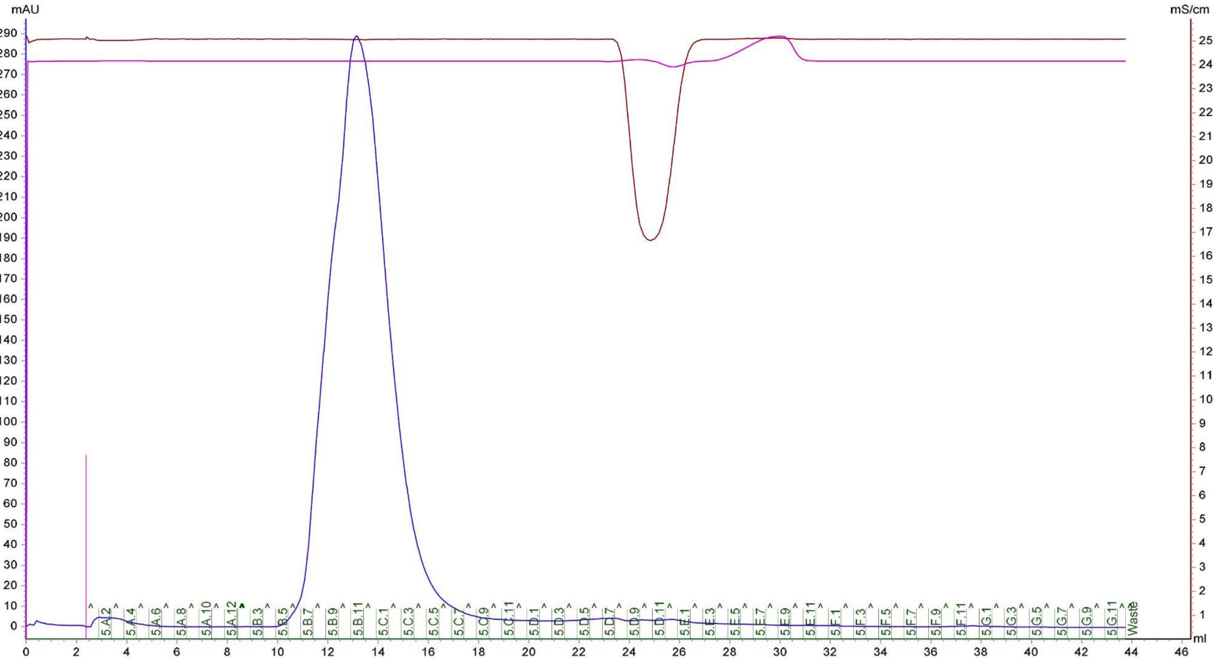


**Supp. Fig. 11.** Gel filtration chromatogram of His-tagged HSA2 variant.

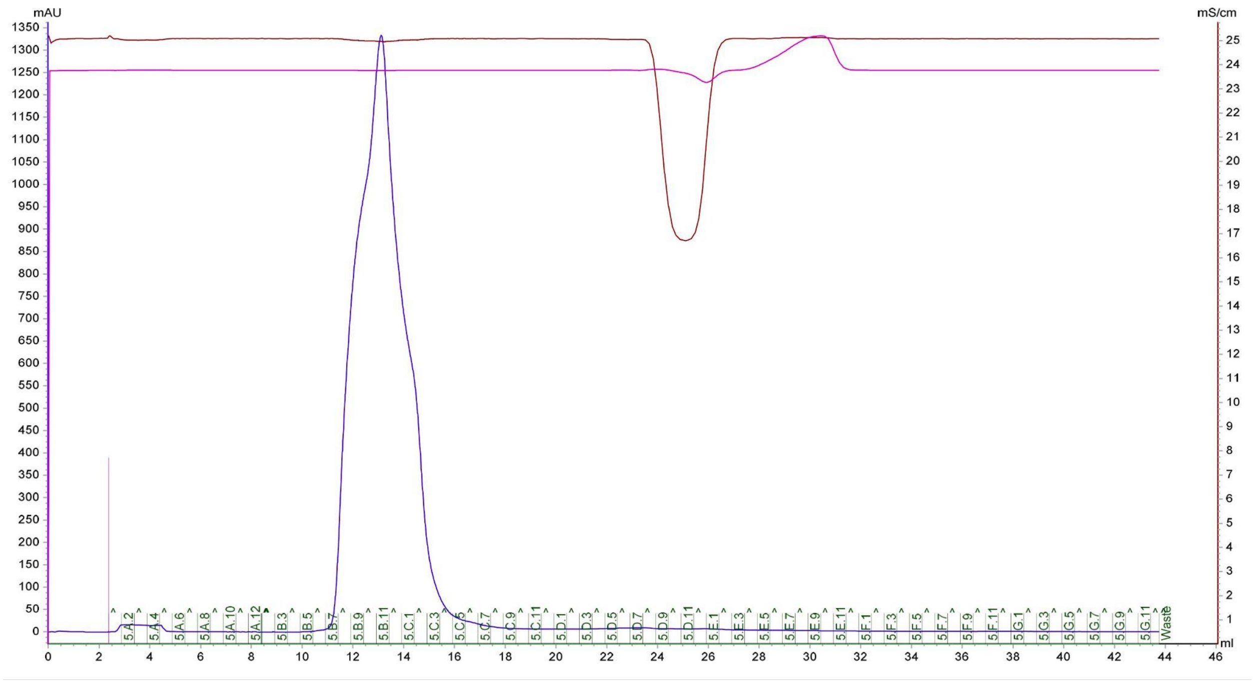


**Supp. Fig. 12.** Gel filtration chromatogram of His-tagged HSA3 variant.

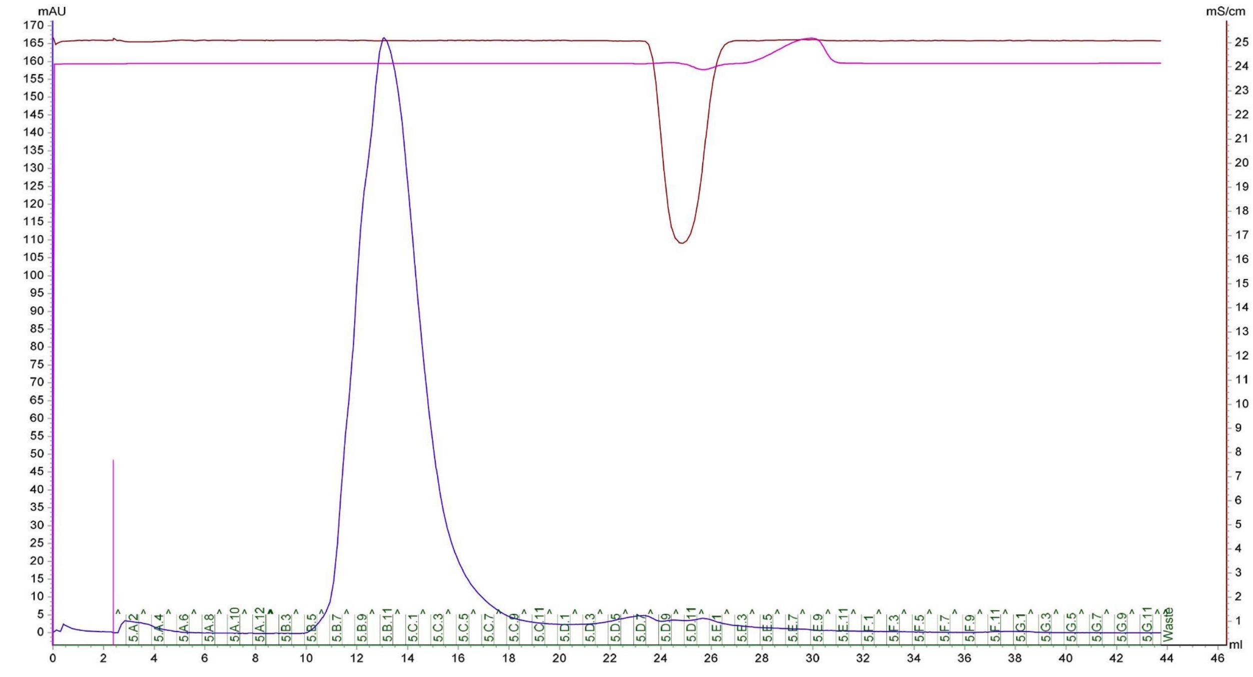


**Supp. Fig. 13.** Gel filtration chromatogram of His-tagged BSA1 variant.

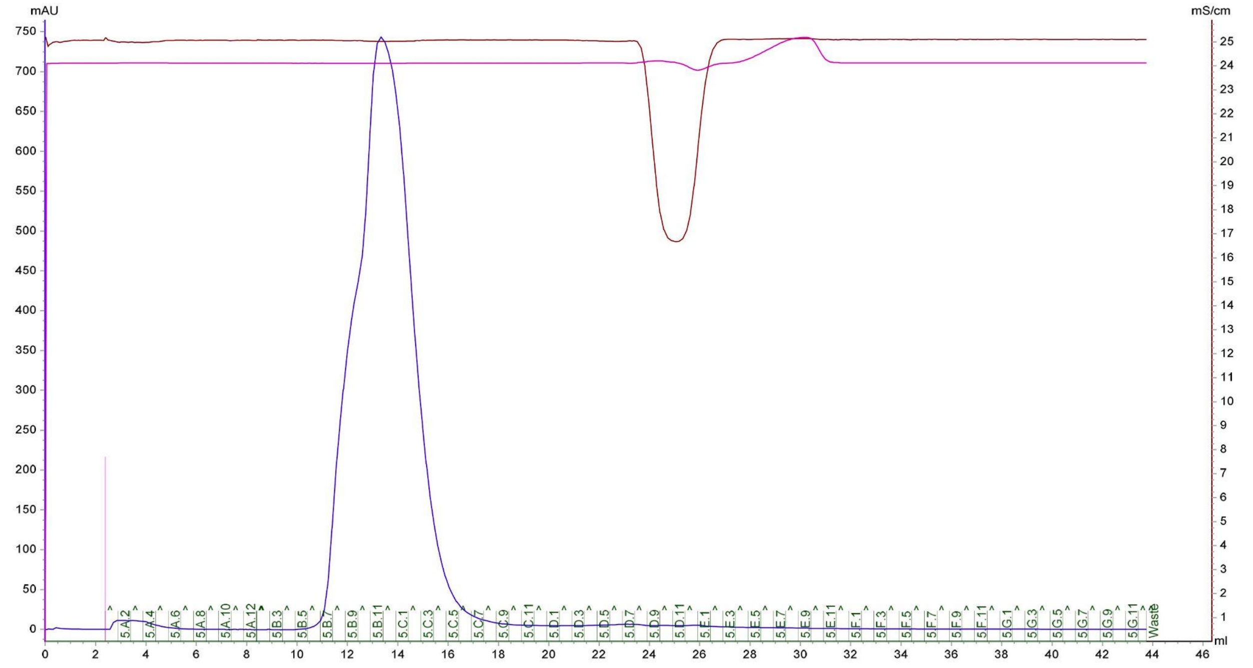


**Supp. Fig. 14.** Gel filtration chromatogram of His-tagged BSA2 variant.

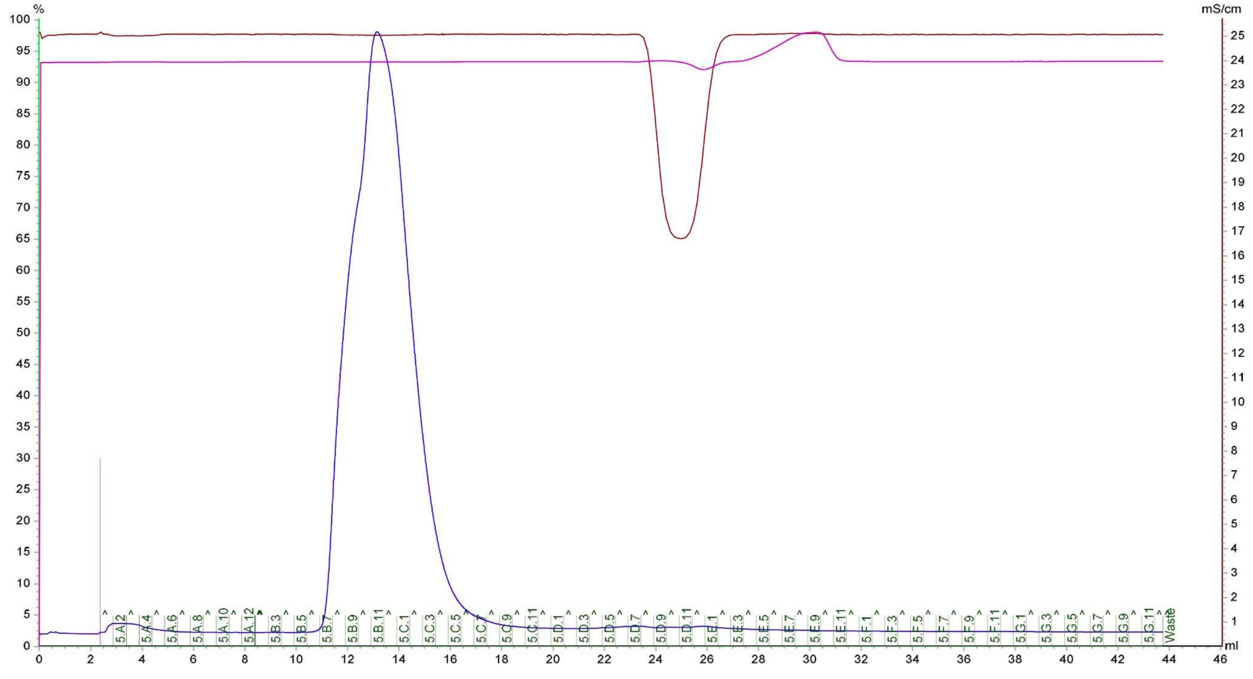


**Supp. Fig. 15.** Gel filtration chromatogram of His-tagged BSA3 variant.

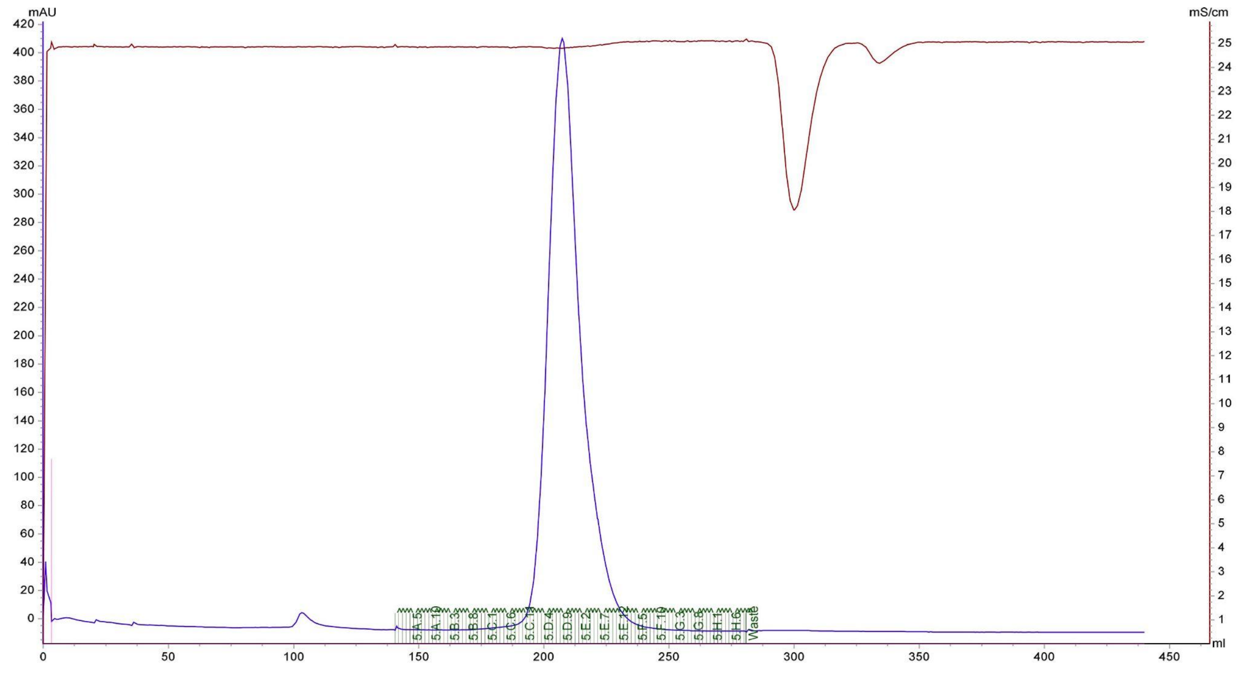


**Supp. Fig. 16.** Gel filtration chromatogram of HSA1-SUMO variant after cleavage of bdSUMO tag.

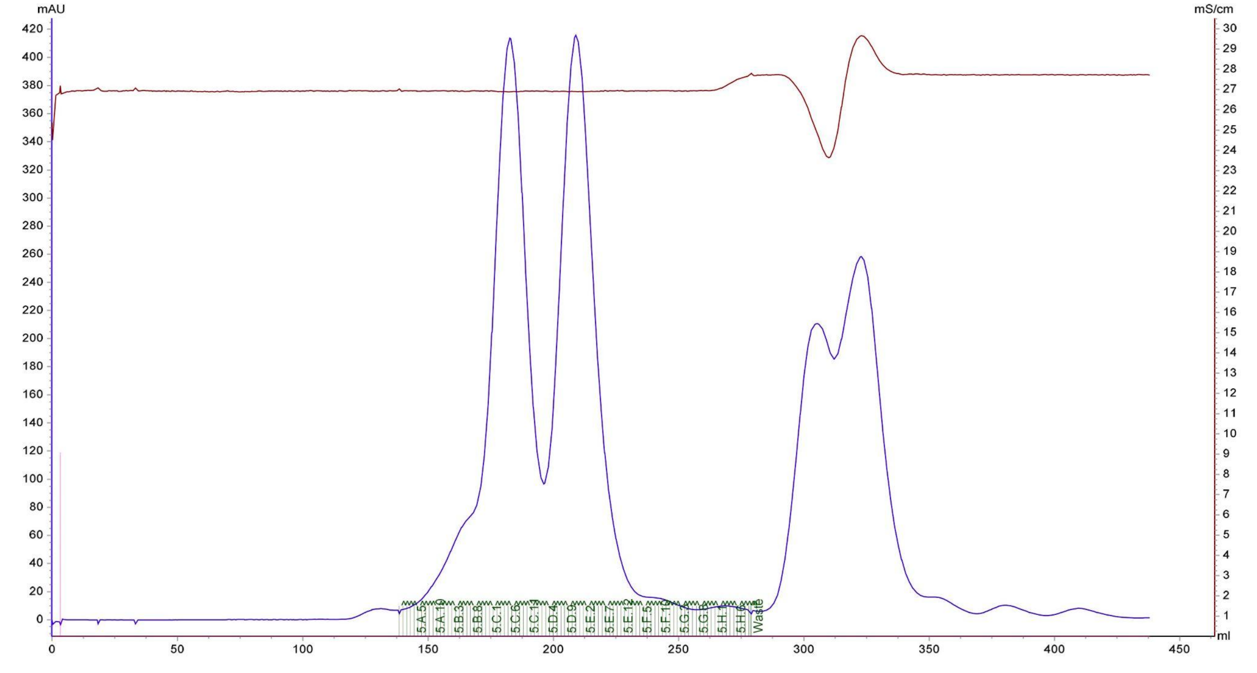


**Supp. Fig. 17.** Gel filtration chromatogram of BSA2-SUMO variant after cleavage of bdSUMO tag. Prep with equal amount of monomeric and dimeric fractions of BSA2.

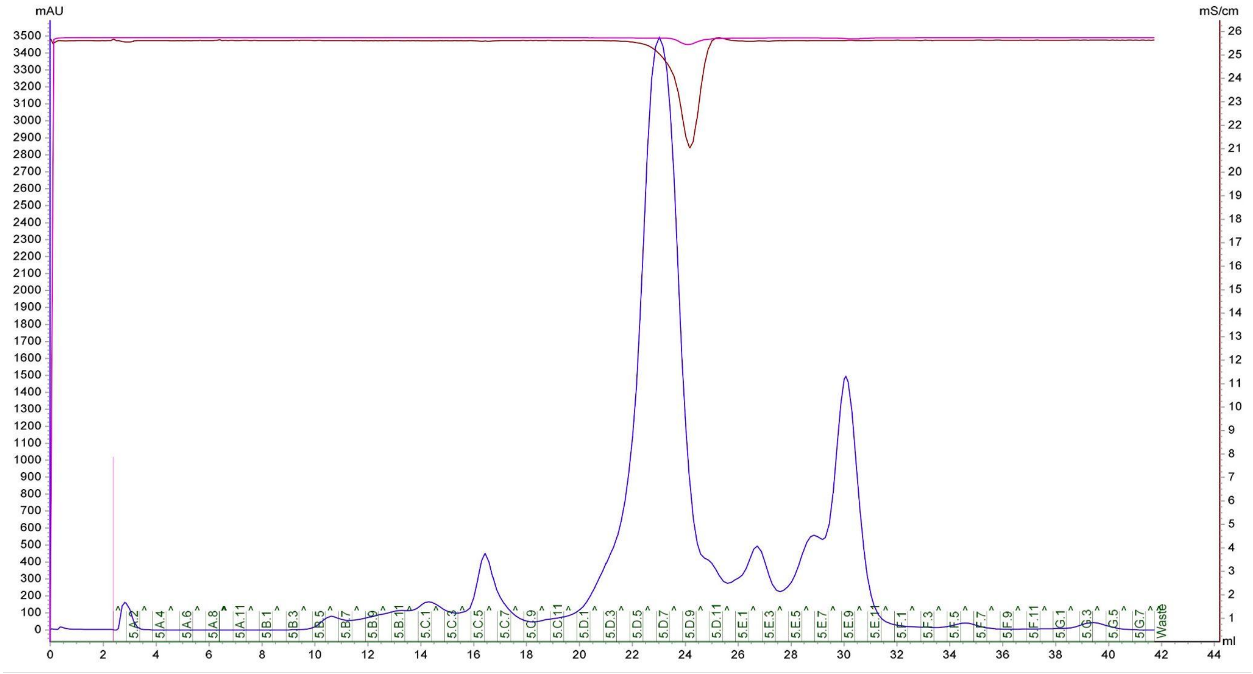


**Supp. Fig. 18.** Gel filtration chromatogram of tagless HSA3 variant.

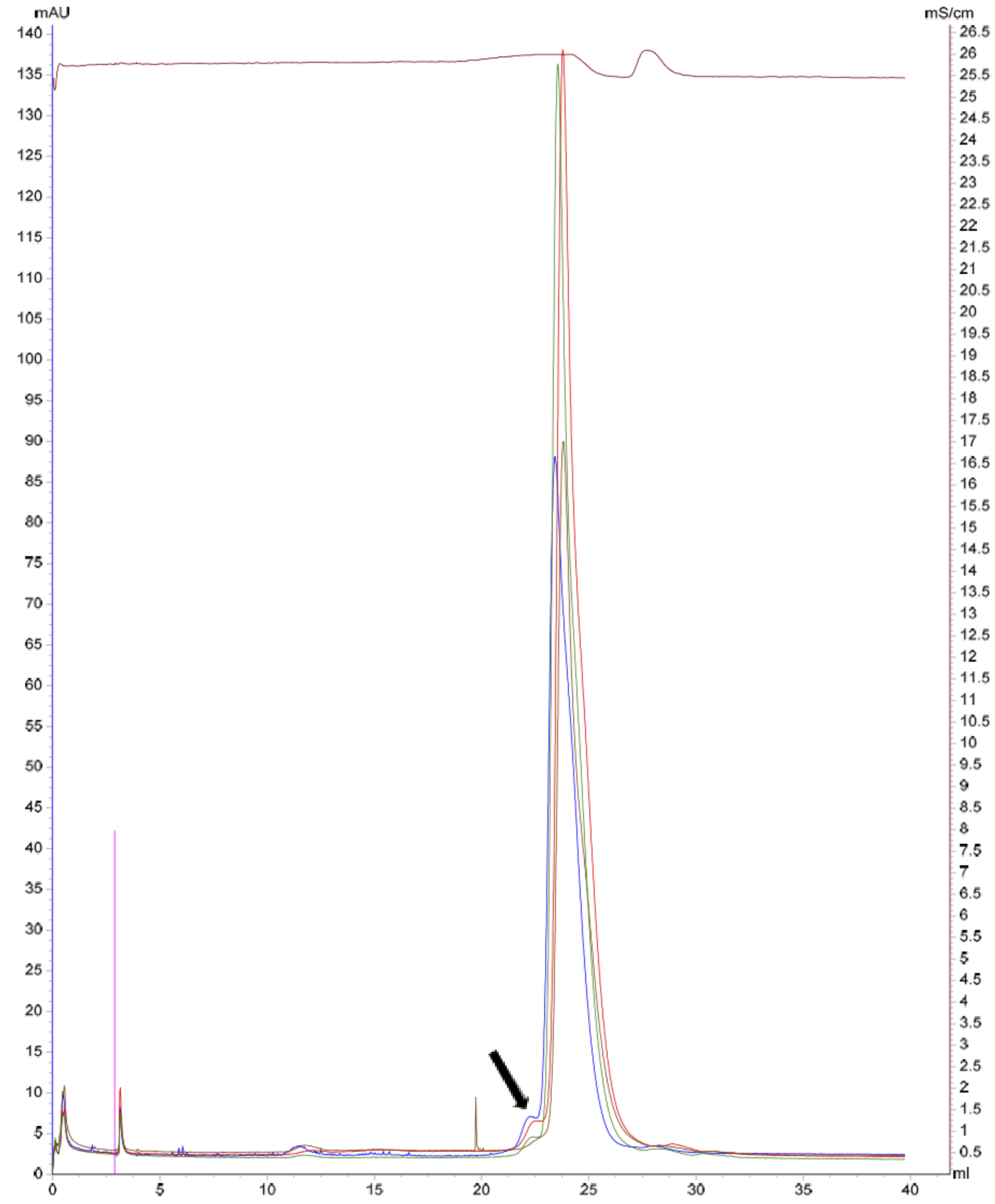


**Supp. Fig. 19.** Analytical gel filtration chromatograms of tagless albumin variants following four months of storage at 4ºC. The chromatograms of BSA2 (green), HSA1 (dark brown), and HSA6 (red) were overlaid with that of a fresh batch of commercial BSA (blue). More than 95% of the proteins were monomeric (main peak). The peaks of the protein dimers is indicated by a black arrow.

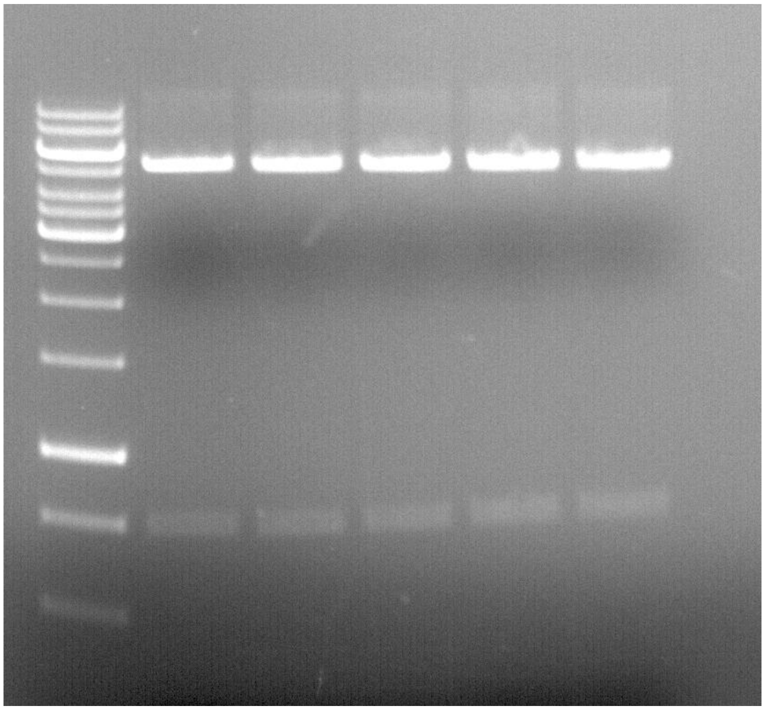

**Supp. Fig. 20.** Restriction of pET29b plasmid with *NcoI* and *XhoI* enzymes, overnight at 37°C. From left to right: DNA ladder, buffer with no albumin, buffer with 0.1mg/ml commercial HSA, buffer with 0.1mg/ml HSA1, buffer with 0.1mg/ml BSA2, buffer with 0.1mg/ml HSA3.
